# Multi-omic rejuvenation and lifespan extension upon exposure to youthful circulation

**DOI:** 10.1101/2021.11.11.468258

**Authors:** Bohan Zhang, David E. Lee, Alexandre Trapp, Alexander Tyshkovskiy, Ake T. Lu, Akshay Bareja, Csaba Kerepesi, Lauren H. Katz, Anastasia V. Shindyapina, Sergey E. Dmitriev, Gurpreet S. Baht, Steve Horvath, Vadim N. Gladyshev, James P. White

## Abstract

Heterochronic parabiosis (HPB) is known for its functional rejuvenation effects across several mouse tissues. However, its impact on the biological age of organisms and their long-term health remains unknown. Here, we performed extended (3-month) HPB, followed by a 2-month detachment period of anastomosed pairs. Old detached mice exhibited improved physiological parameters and lived longer than control isochronic mice. HPB drastically reduced the biological age of blood and liver based on epigenetic analyses across several clock models on two independent platforms; remarkably, this rejuvenation effect persisted even after 2 months of detachment. Transcriptomic and epigenomic profiles of anastomosed mice showed an intermediate phenotype between old and young, suggesting a comprehensive multi-omic rejuvenation effect. In addition, old HPB mice showed transcriptome changes opposite to aging, but akin to several lifespan-extending interventions. Altogether, we reveal that long-term HPB can decrease the biological age of mice, in part through long-lasting epigenetic and transcriptome remodeling, culminating in the extension of lifespan and healthspan.

## INTRODUCTION

Aging is the primary risk factor for chronic diseases (Brett and Rando, 2014; Lopez-Otin et al., 2013). It brings accumulation of damage at many levels of biological organization and a pervasive and destructive decline of organ function, resulting in inevitable mortality. Although many attempts have been made to extend lifespan and ameliorate specific aging phenotypes through interventions, aging itself has generally been regarded as an irreversible process. Currently, there is no clear evidence that any intervention can rewind the biological age of a whole organism. With recent developments of advanced aging biomarkers based on DNA methylation (i.e., methylation clocks) (Horvath, 2013; Meer et al., 2018; Olova et al., 2019; Petkovich et al., 2017), the concept of aging as an irreversible process has been challenged via precise measurements of biological age. With their convincing assessment of the attenuated aging effect of various longevity interventions, including caloric restriction (CR), genetic models and reprogramming, methylation clocks have been generally recognized as a robust readout of organismal biological age. Notably, DNA methylation clocks have successfully predicted the reversal of biological age by several interventions, including reprogramming factor expression and treatment with drugs or blood components (Fahy et al., 2019; Horvath et al., 2020; Lu et al., 2020; Rando and Chang, 2012; Sarkar et al., 2020). Clocks have been also recently used to reveal and describe a natural rejuvenation event occurring during early embryonic development (Kerepesi et al., 2021; Trapp et al., 2021). However, in the case of interventions, it remains generally enigmatic whether the predicted reversal of biological age is sustained, correlated with longer lifespan, and manifests in improved physiological function.

The heterochronic parabiosis (HPB) model has been used to study circulating factors that regulate the aging process since the 1950s (Lunsford et al., 1963; McCay et al., 1956; Pope et al., 1956). More recent work has established the model as a proof of concept that youthful circulation can restore old tissue functions (Baht et al., 2015; Conboy et al., 2005). Indeed, the effects of HPB on the amelioration of aging phenotypes are evident across tissues including muscle (Conboy et al., 2005), liver (Conboy et al., 2005), heart (Loffredo et al., 2013), brain (Ruckh et al., 2012; Villeda et al., 2014) and bone (Baht et al., 2015; Vi et al., 2018). Remarkably, these effects are observed typically after only 4-5 weeks of parabiosis. Similar results are observed with acute heterochronic blood exchange (non-parabiosis), showing beneficial effects of “young blood” on muscle, liver and brain of old recipients (Rebo et al., 2016). Heterochronic young blood plasma transfer also improves pathology of age-related diseases, such as in a model of Alzheimer’s disease in mice (Middeldorp et al., 2016). Although HPB leads to diverse effects on old cells and tissues, our understanding of the molecular mechanisms involved remains limited. Likewise, whether the immediate effects of blood/plasma exchange are sustained after the procedure is still unknown. Lastly, due to the previous absence of a precise quantification method, it remains unclear whether HPB can slow or rewind the biological aging of organisms.

A previously reported (Conboy et al., 2013; Wright et al., 2001), but seldomly used HPB procedure is the detachment of mice following parabiosis. This technique has previously been used to verify cell engraftment using different tracer techniques (Donskoy and Goldschneider, 1992). A recent investigation revealed the persistence of aged hematopoietic stem cells in the young bone marrow niche resulting from HPB months after a surgical separation of parabionts (Ho et al., 2021); however, to our knowledge there have been no studies to investigate longevity or long-term effects on healthspan. Although detachment involves performing a second surgery on the animals, it permits physiological and longevity measurements, which are difficult to obtain in the case of anastomosed mice.

Here, we report the results of a long-term HPB study followed by a detachment period. We found that old mice detached from young mice showed an extended lifespan and improvements across several dimensions of aging biology. By comprehensive epigenetic clock and RNA-seq analyses, we observed a robust reduction in biological age of old mice following 3-month HPB, sustained even after a detachment period of two months. Notably, this rejuvenation effect was significantly stronger than that observed upon short-term HPB. We find that transcriptomic and epigenetic profiles of long-term HPB are intermediate between young and old isochronic pairs, and that HPB positively associates with the effects of common lifespan-extending interventions and counteracts aging-related gene expression changes. Our findings suggest the presence of profound and persisting molecular rejuvenation effects following exposure to young circulation, leading to extended lifespan and healthspan.

## RESULTS

### Long-term parabiosis followed by detachment extends lifespan and healthspan in mice

We used a long-term (3-month) parabiosis (either heterochronic or isochronic) period in mice, followed by detachment of the parabionts. Young mice started parabiosis at 3 months and old mice at 20 months of age **(Figure 1A)**. Transcriptomic and epigenetic profiling of the liver and blood was conducted to assess molecular changes resulting from HPB **(Figure 1B)**. For longevity and healthspan experiments, mice were detached and allowed 1 month of recovery after parabiosis prior to physiological data collection **(Figure S1A)**. After separation from respective parabiosis pairs, some mice were allowed to live freely until their natural death, in order to examine the effect of parabiosis on lifespan. We observed a significant 6-week extension in median lifespan and a 2-week extension in maximum lifespan in old mice detached from heterochronic pairs as compared to their isochronic controls **(Figure 1C)**. This was accompanied by an initial reduction in body weight coupled with better preservation of body weight in the final months of life **(Figure 1D)**. The initial drop in body weight appeared to be primarily caused by a reduction in fat mass, while the preservation of body weight was due to the maintenance of both lean and fat mass later in life **(Figure 1D)**. These changes in body composition were independent of changes in food consumption **(Figure S1B)**. In addition to improvements in body composition, old heterochronic mice showed higher voluntary cage activity than isochronic controls **(Figure S1C)**. While it is possible that old heterochronic mice received some small training effect due to attachment to young, more active partners, mouse physical capacity decreases rapidly following aerobic exercise training, returning to pre-training levels by 4 weeks (Brace et al., 2016). This detraining would be apparent by the second month following detachment, if the training were the sole contributor to the increased physical activity; however, the observed increases in cage activity persist for at least 7-9 months **(Figure 1D, Figure S1C)**.

**Figure 1.**
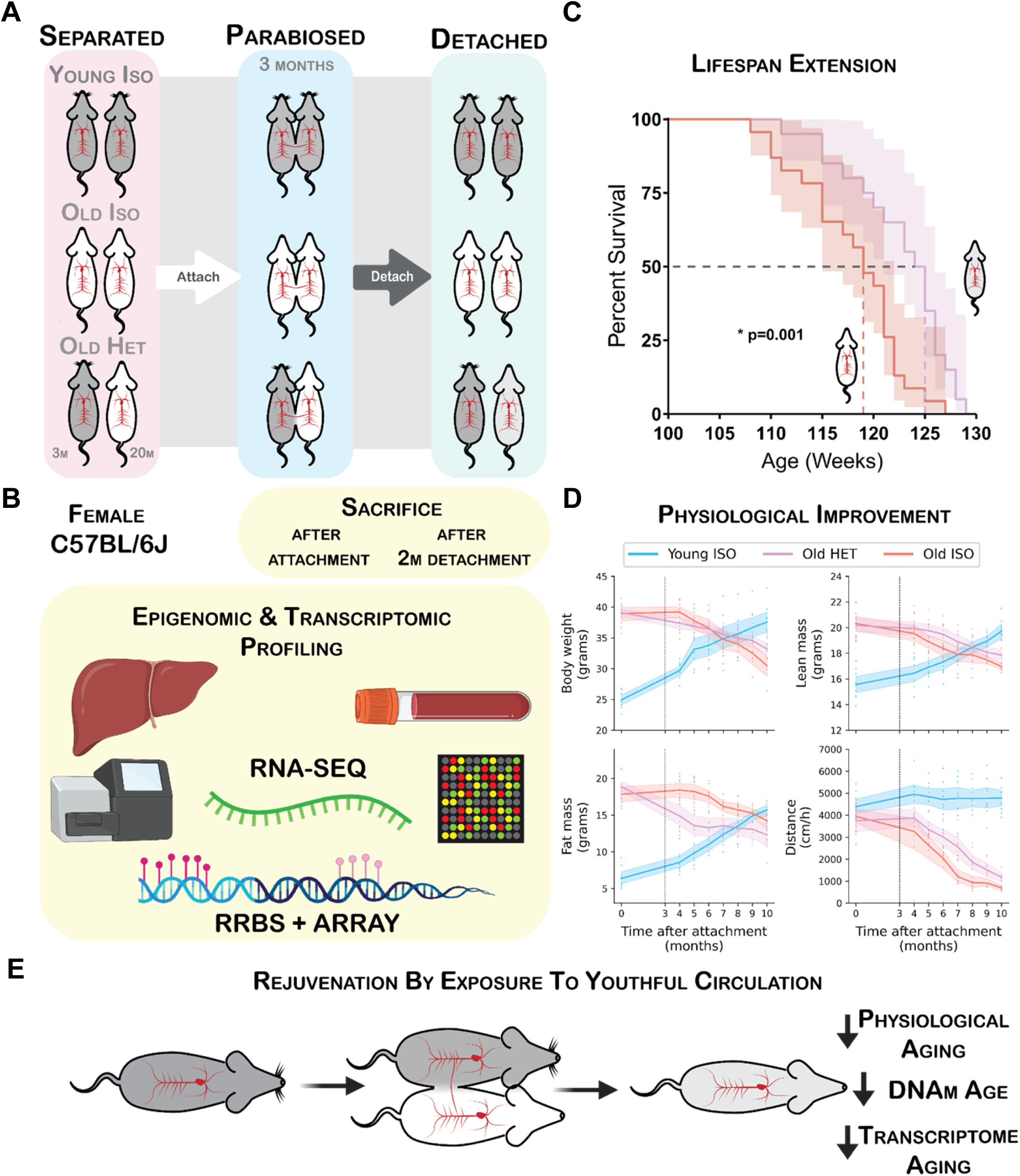
Prolonged parabiosis followed by detachment leads to extended lifespan and healthspan. **(A)** Overview of the parabiosis and detachment model. All pairs were anastomosed for 3 months, starting at 20 months of age for old mice and 3 months of age for young mice. At 23 months, old mice were detached, and remaining lifespan was assessed. **(B)** Schematic of the molecular profiling methods applied to mouse tissues. **(C)** Survival curve of detached old mice from isochronic (red, *n* = 23) and heterochronic (purple, *n* = 20) pairs. Kaplan-Meier curves with 95% confidence interval are shown. Kaplan-Meier survival analysis was used for statistical testing. **(D)** Body weight, lean mass, fat mass and cage activity measurements before parabiosis and at monthly time points starting 1 month after detachment. Mean values with 95% confidence intervals are shown. Statistical analysis (described in the Methods) was carried out between old isochronic and heterochronic groups to determine the significance of the difference: p(body weight) = 0.29, p(lean mass) = 0.049, p(fat mass) = 0.00092, and p(cage activity) = 0.015. Young ISO denotes young mice from isochronic pairs, Old ISO denotes old mice from isochronic pairs, Old HET denotes old mice from heterochronic pairs. Dashed lines show the time of detachment. *n* = 7-10/group. **(E)** Schematic of the outlined results of the study, wherein long-term parabiosis leads to sustained epigenomic, transcriptomic, and physiological rejuvenation.

### Epigenetic age of old mice is reversed by parabiosis with a sustained effect after detachment

For epigenetic analyses, we subjected mice to the same prolonged attachment, and harvested tissues either immediately after the 3-month parabiosis procedure, or 2 months after detachment **(Figure S1A)**. We first subjected the blood and liver of the animals to Reduced Representation Bisulfite Sequencing (RRBS)(Meissner et al., 2005) **(Figure 2A, Table S1)**. Since blood samples taken immediately after detachment contained a mixture of old and young blood **(Figure S2)**, we initially chose to quantify methylation changes in blood after 2-month detachment to investigate whether epigenomic remodeling upon exposure to young circulation persists without young blood contamination. Additionally, we took liver as an example of a solid tissue to quantify the indirect effects on the methylome happening immediately after parabiosis and following 2-month detachment.

**Figure 2.**
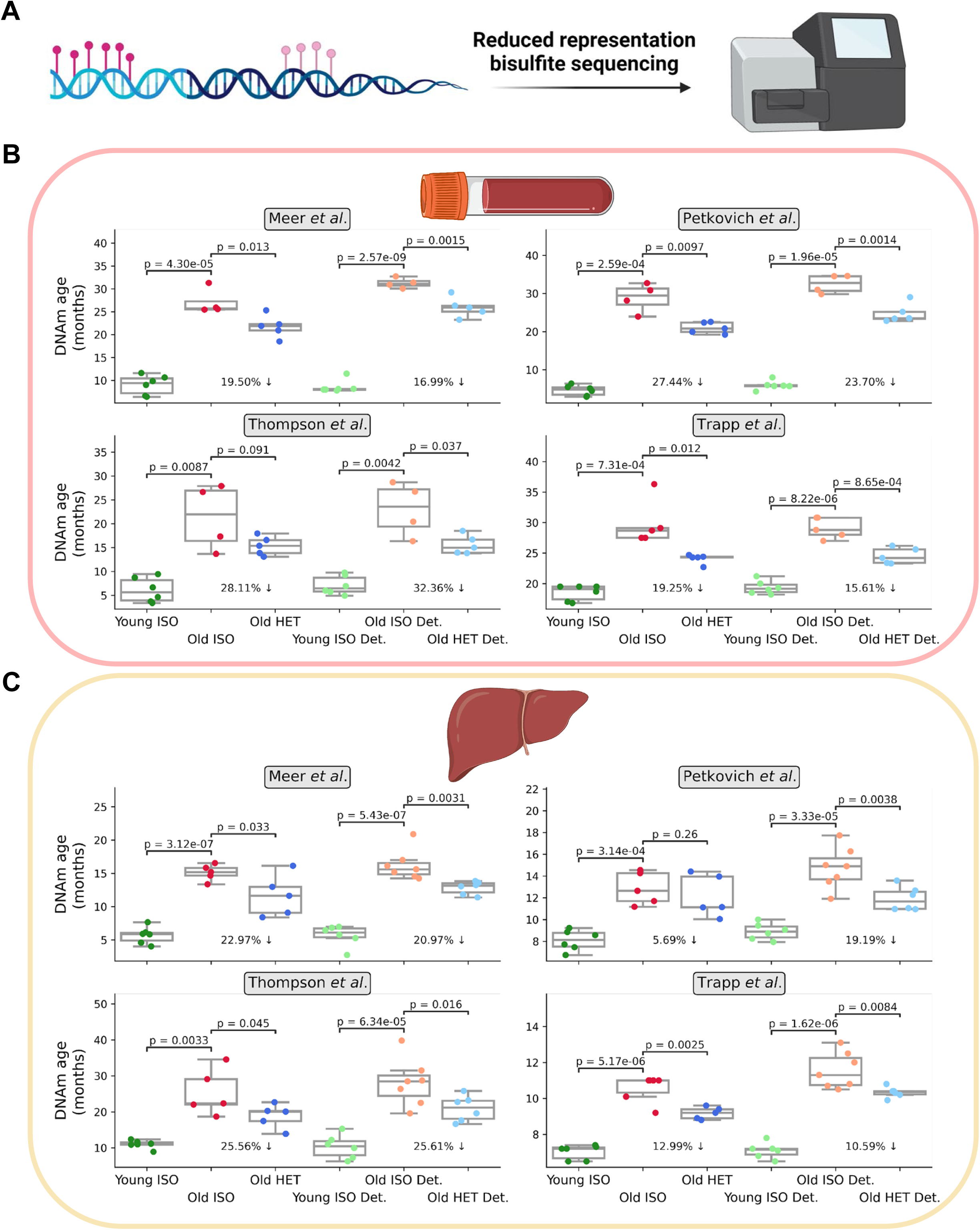
Persistent epigenetic age reversal in blood and liver upon HPB assessed by RRBS-based aging clocks. **(A)** Schematic of the experiment. DNA cytosine methylation was assessed by high-depth reduced representation bisulfite sequencing. **(B)** DNA methylation age of old heterochronic blood before and after detachment plotted with isochronic young and old controls assessed by the Meer et al., Petkovich et al., Thompson et al., and Trapp et al. clocks. *n* = 4-6 per group. **(C)** DNA methylation age of old heterochronic liver before and after detachment plotted with isochronic young and old controls assessed by the Meer et al., Petkovich et al., Thompson et al., and Trapp et al. clocks. *n* = 5-7 per group. “Young ISO” denotes young mice from isochronic pairs, “Old ISO” denotes old mice from isochronic pairs, and “Old HET” denotes old mice from heterochronic pairs. “Det.” denotes detached old mice from isochronic or heterochronic pairs. Epigenetic ages are shown in months. One-tailed Welch’s t-tests assuming unequal variances were used for statistical analyses.

To quantify the biological age of the animals, we applied four RRBS-based epigenetic clocks to 6 different groups of mice: young isochronic, old isochronic, and old heterochronic mice, as well as detached variants of all these three groups. Clocks used included two recently developed multi-tissue clocks (Meer et al., 2018; Thompson et al., 2018), a blood-specific clock (Petkovich et al., 2017), and the first single-cell clock framework (Trapp et al., 2021), which was modified to accommodate and profile epigenetic age in bulk data. Application of these clocks to blood RRBS data revealed a profound epigenetic age decrease (19-28%) when comparing old isochronic and heterochronic pairs. Remarkably, this effect was sustained even after two months of detachment (reduction in age by 16-32%) **(Figure 2B)**. Application of these same four clocks to liver data showed comparable immediate (5-26%) and sustained (10-26%) epigenetic age reduction as observed in blood **(Figure 2C)**. Together, these results indicate that long-term HPB rejuvenates the epigenome compared to isochronic pairs in both blood and liver. Moreover, these unbiased rejuvenation signals in solid organs (such as the liver) further suggest that long-term HPB acts in a systemic manner, leading to global epigenomic remodeling and organism-wide age reversal.

To confirm this biological age decrease, we employed the use of a completely independent epigenomics analysis platform: methylation profiling of liver samples on the recently developed Mammalian Methylation Array (Lu et al., 2021) **(Figure 3A, Table S2)**. We performed this profiling across 3 different long-term parabiosis groups (young isochronic, old isochronic, and old heterochronic) along with detached pairs of each of these groups. We applied four different array-specific epigenetic clocks. Two universal clocks were used: one based on relative age standardized to maximum lifespan and one based on log-linear transformed age standardized to the age of sexual maturity. Additionally, two mouse-specific clocks were used: a broad liver-specific clock and a clock tracking the epigenetic dynamics of the developmental process in the liver. Together, all four clocks were able to correctly predict the age of control isochronic samples with high precision and accuracy, and showed a highly significant epigenetic age decrease when comparing old heterochronic and isochronic mice (17-27% reduction in age) **(Figure 3B)**. Remarkably, this effect was preserved after detachment, with a slight reduction in effect size (11-21% reduction in age). Moreover, results from different platforms and organs were well correlated with each other (**Figure S3**). These results from an independent platform further reinforce the notion that 3-month HPB leads to a profound, sustained, and systemic epigenetic rejuvenation.

**Figure 3.**
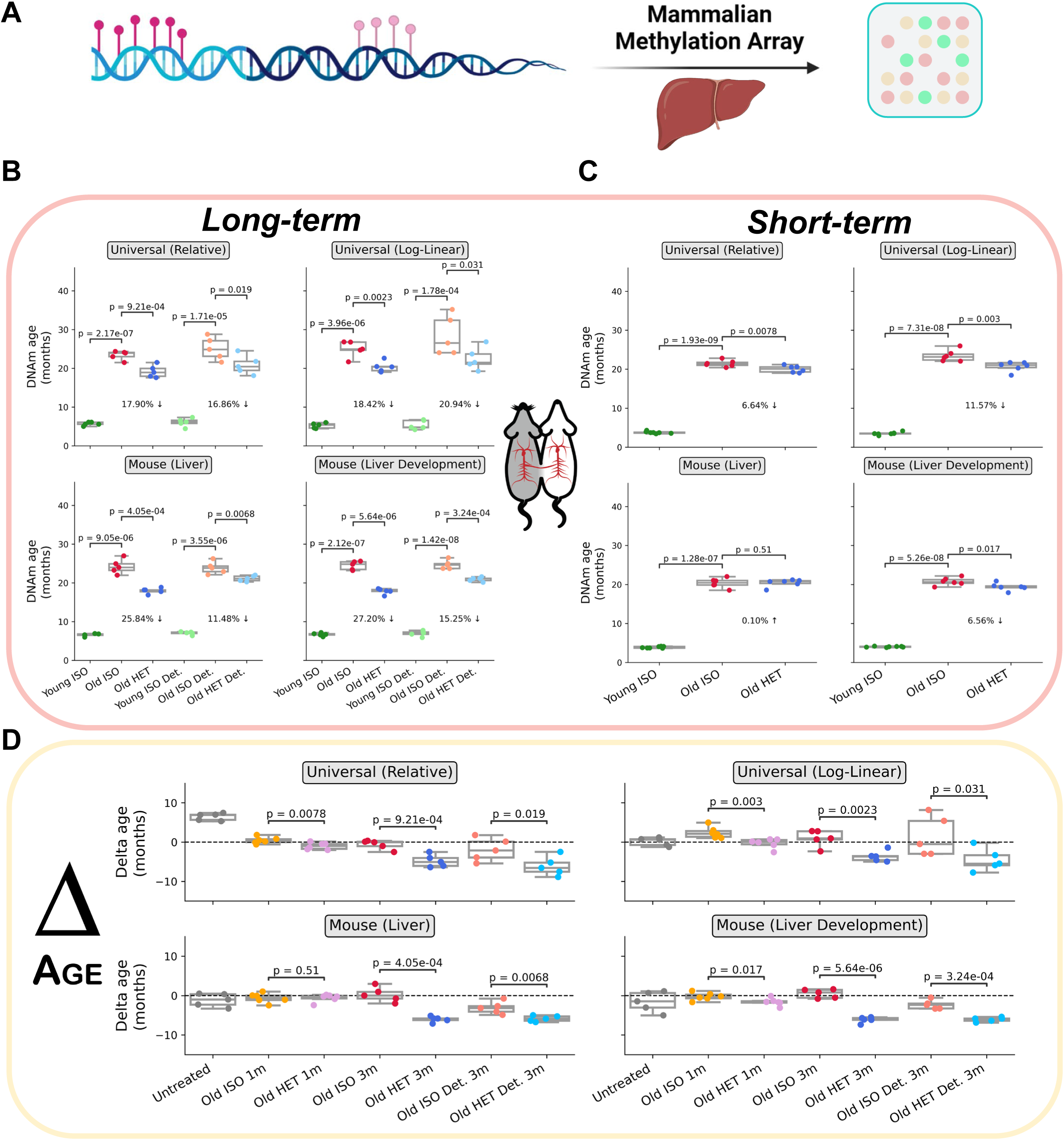
Sustained epigenetic age reversal in liver upon HPB assessed by microarray-based aging clocks. **(A)** Schematic of the experiment. DNA methylation was assessed via the recently developed mammalian methylation array (HorvathMammalMethylChip40). **(B)** Epigenetic age of long-term (3-month) parabiosis liver samples from old heterochronic attached or detached mice, plotted with young and old isochronic controls, based on the universal relative age mammalian, universal log-linear transformed age mammalian, liver, and development liver clocks. *n* = 5 per group. **(C)** Epigenetic age of short-term (5-week) parabiosis liver samples from old heterochronic attached or detached mice, plotted with young and old isochronic controls, based on the universal relative age mammalian, universal log-linear transformed age mammalian, liver, and development liver clocks. *n* = 6-7 per group. **(D)** Delta age (epigenetic age minus chronological age) of liver samples from old heterochronic mice, old isochronic controls and untreated controls based on universal relative age mammalian, universal log-linear transformed age mammalian, liver, and development liver clocks. Dashed line denotes a delta age of 0 (epigenetic age = chronological age). Points above this line depict age acceleration (epigenetic age > chronological age), while points below this line depict age deceleration (epigenetic age < chronological age). *n* = 5 per group. One-tailed Welch’s t-tests assuming unequal variances were used for statistical analyses.

Furthermore, to test the difference between the commonly used 5-week short-term protocol and our long-term (3-month) HPB protocol, methylation array profiling of liver tissue was also performed in three groups of animals (young isochronic, old isochronic, and old heterochronic) immediately after a 5- week attachment period. Only three of the four clocks applied detected a significant reduction in epigenetic age in heterochronic mice, and the effect size was dramatically smaller than in the long-term (3-month) parabiosis experiment (only 0-11% reduction in age with short term HPB) **(Figure 3C)**. Notably, the clock trained specifically on mouse liver tissues did not detect this reduction. This suggests that while short-term parabiosis may induce some moderate epigenetic remodeling leading to mild rejuvenation readouts, a longer-term treatment clearly outperforms short-term attachment when it comes to immediate and particularly sustained rejuvenation of the liver epigenome.

To further determine whether an age-reversal (i.e., rejuvenation) occurs, an additional analysis of delta ages (the signed difference between epigenetic and chronological age) across all groups, including an untreated control group, was performed **(Figure 3D)**. This revealed that heterochronic pairs, both after 3-month attachment and after the 2-month detachment period, exhibited significantly lower epigenetic age compared to their chronological age. The same trend was observed with attached mice from short-term parabiosis experiments, but the effect size was dramatically stronger upon long-term treatment. Additionally, we found that mean methylation was consistent among samples, suggesting that sustained, targeted epigenetic remodeling, manifesting itself in decreased epigenetic clock age, occurs as a result of long-term HPB (**Figure S4**).

Taken together, we report that a profound and systemic rejuvenation occurs in old animals upon exposure to youthful circulation, as assessed by 8 distinct epigenetic clocks across two independent DNA methylation analysis platforms. We further find that this age-reversal phenotype, driven by epigenetic remodeling, is sustained even after a 2-month detachment period. These results present the first example of a systemic *in vivo* rejuvenation quantified by molecular biomarkers in mice.

### Gene expression analyses reveal pathways responsive to long-term parabiosis

Given the remarkable epigenetic age reversal, we sought to concurrently elucidate the transcriptomic changes that occur as a result of HPB. We performed RNA-seq analyses of liver tissues in old isochronic and heterochronic mice, as well as in detached groups of both conditions, followed by differential gene expression and gene set enrichment analyses (GSEA) **(Figure 4A)**. Using the Hallmark, KEGG, REACTOME and GO BP gene sets obtained from MSigDB (Liberzon et al., 2015; Subramanian et al., 2005), we computed normalized enrichment scores comparing isochronic and heterochronic samples immediately after the 3-month parabiosis period, as well as after 2-month detachment **(Figure 4B, 6E)**. Among samples taken immediately following the attachment period, we observed strong positive enrichment in the oxidative phosphorylation and mitochondrial biogenesis gene sets. Interestingly, oxidative phosphorylation is known to be disrupted during the aging process (Lesnefsky and Hoppel, 2006). Within this context, our results indicate that HPB may reverse some of the age-associated decline commonly associated with this critical energy production pathway. We also observed negative enrichment of inflammatory and interferon gamma (IFN-gamma) response gene sets. IFN-gamma protein level is known to increase with age in some tissues (particularly in T-cells) (Bandres et al., 2000), and inflammation is one of the crucial hallmarks of aging (Lopez-Otin et al., 2013) associated with the development of various age-related pathologies, including cancer (Leonardi et al., 2018), Alzheimer’s disease (Lai et al., 2017) and chronic kidney disease (Amdur et al., 2016). These results further support the rejuvenating effect of heterochronic parabiosis on murine transcriptomic profiles.

**Figure 4.**
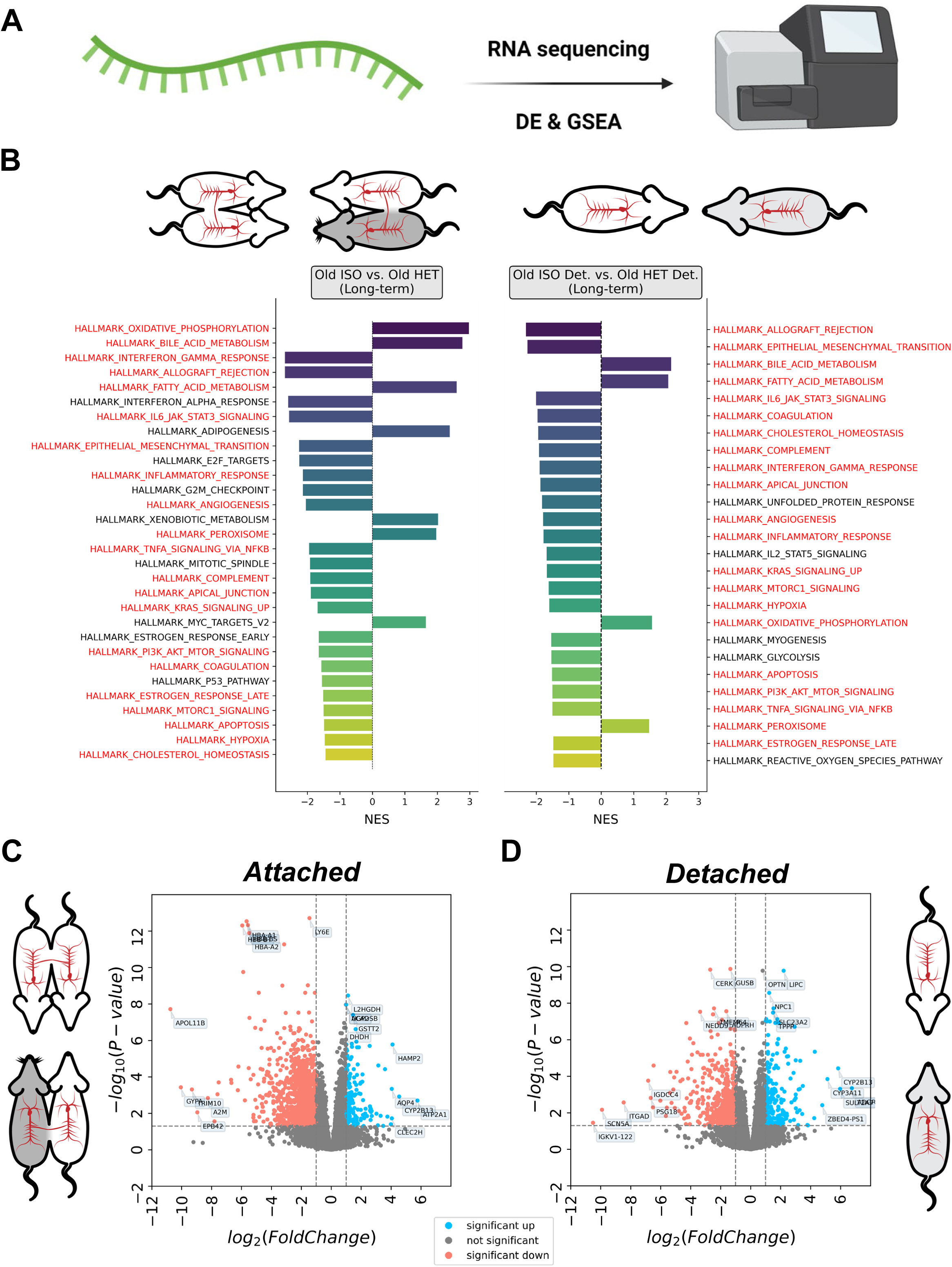
Transcriptomic analyses of HPB reveal unique pathway enrichment and differential expression patterns. **(A)** Schematic outline of the analyses. RNA was isolated from liver tissue of old heterochronic and isochronic mice, either immediately after detachment or after a 2-month detachment period. Differential expression (DE) and gene set enrichment analyses (GSEA) were subsequently performed on the resulting transcriptomic data. **(B)** GSEA of old isochronic vs old. heterochronic mice (left), as well as of detached old isochronic vs. detached old heterochronic mice (right). Hallmark gene sets (n = 50) were used as input to GSEA. Only significant enrichments (adjusted p-value < 0.05) are shown. Gene sets shown in red are commonly enriched in the same direction in both the attached GSEA and detached GSEA. Significant gene sets are ranked from top to bottom based on the absolute value of the normalized enrichment score. **(C-D)** Volcano plot of differentially expressed genes in attached **(C)** and detached **(D)** old heterochronic (*n* = 3) and old isochronic (*n* = 3) mice. Some highly significant genes are shown in each plot. Genes shown in red are significantly downregulated in heterochronic mice, while genes shown in blue are significantly upregulated in heterochronic mice, after multiple testing correction.

**Figure 5.**
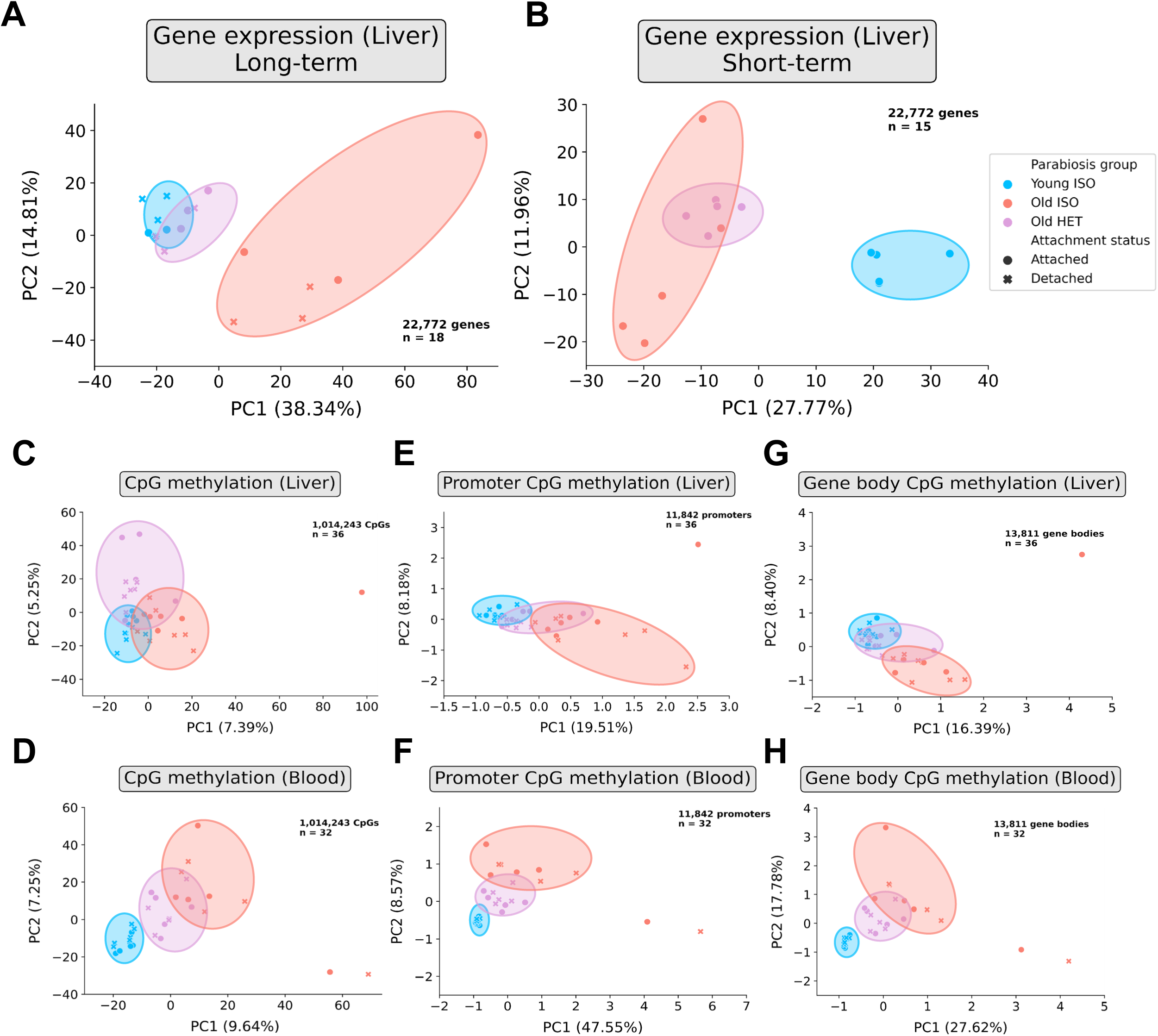
Dimensionality reduction highlights rejuvenated molecular profiles following HPB. **(A)** Principal Component Analysis (PCA) of liver RNA-seq data of long-term parabiotic mice or 2 months after detachment (*n* = 3 per group). **(B)** PCA of liver RNA-seq data of short-term parabiotic mice (*n* = 3 per group) **(C)** PCA of liver RRBS data across all highly covered common CpGs (1,014,243 CpGs, *n* = 5-7 per group) **(D)** PCA of blood RRBS data across all highly covered common CpGs (1,014,243 CpGs, *n* = 5-6 per group) **(E)** PCA of liver RRBS data across gene promoters (11,842 promoters, *n* = 5-7 per group) **(F)** PCA of blood RRBS data across gene promoters (11,842 promoters, *n* = 5-6 per group) **(G)** PCA of liver RRBS data across gene bodies (13,811 gene bodies, *n* = 5-7 per group) **(H)** PCA of blood RRBS data across gene bodies (13,811 gene bodies, *n* = 5-6 group) “Young ISO” denotes young isochronic mice, “Old ISO” denotes old isochronic mice, and “Old HET” denotes old heterochronic mice. Attached refers to samples taken immediately after the parabiosis period, while detached refers to samples taken 2 months of detachment. Percentage of variance explained by the first two principal components are shown in parentheses.

**Figure 6.**
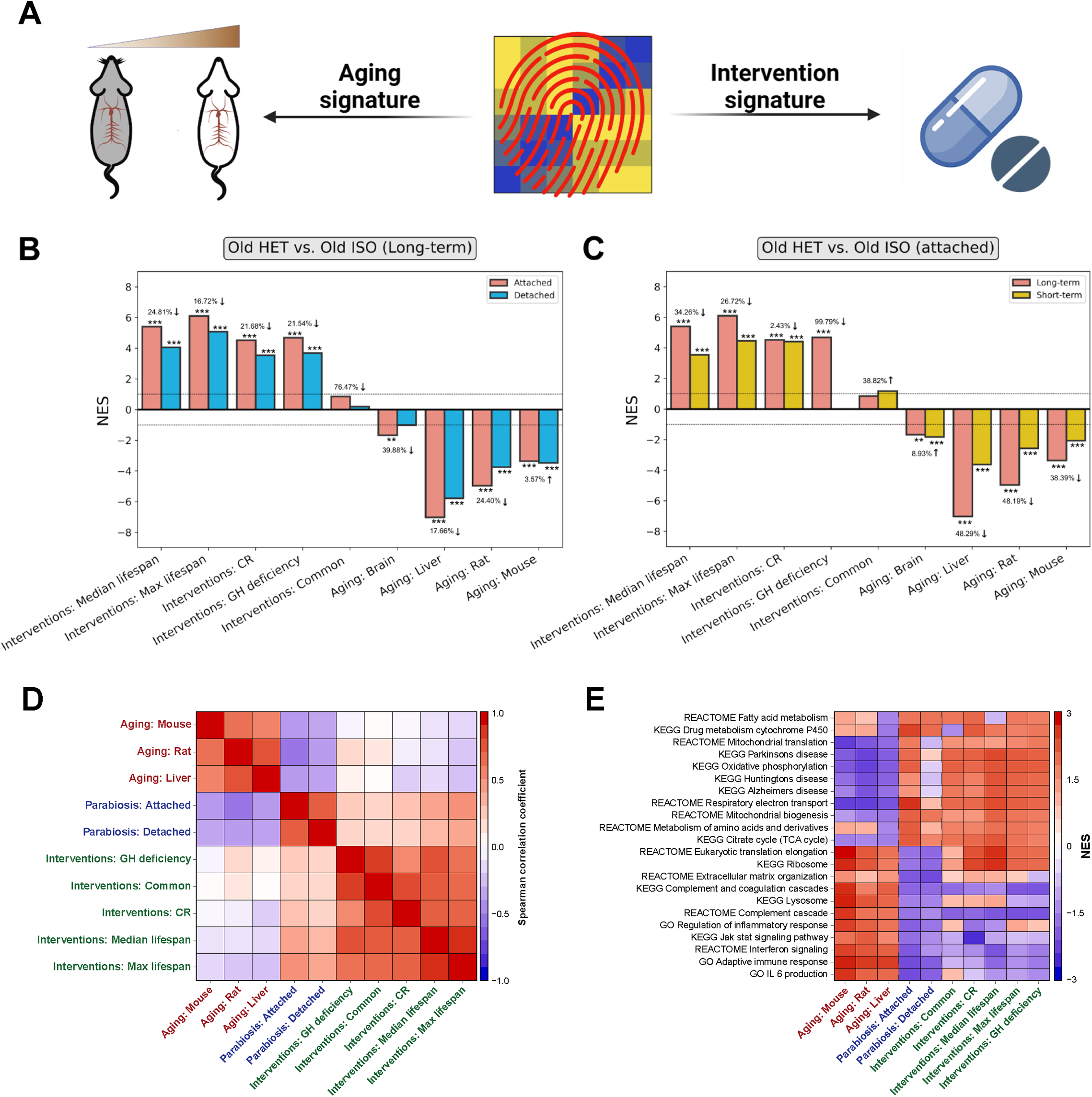
Transcriptomic signature of HPB aligns with longevity interventions and opposes aging. **(A)** Schematic of liver RNA-seq signature analyses. RNA-seq signatures of parabiosis are compared with aging and longevity intervention signatures, following the protocol outlined in (Tyshkovskiy et al., 2019). **(B)** Association between gene expression changes induced by long-term parabiosis with (red) & without detachment (blue) and signatures of aging and lifespan extension. The latter include gene signatures of individual interventions (caloric restriction and growth hormone deficiency), common interventions signatures (Interventions: common) and signatures associated with an effect on lifespan (maximum and median lifespan). The percent decrease comparing attached and detached sample is shown for each pair of bars. * p.adjusted < 0.05; ** p.adjusted < 0.01; *** p.adjusted < 0.001. **(C)** Association between gene expression changes induced by long-term (red) and short-term (yellow) parabiosis without detachment and signatures of aging and lifespan extension. The signatures analyzed are the same as in (B). **(D)** Spearman correlation matrix of gene expression signatures associated with aging (red labels), lifespan extension (green labels) and heterochronic parabiosis (blue labels). **(E)** Functional enrichment analyses of gene expression signatures. Only functions significantly associated with at least one signature are shown. Cells are colored based on normalized enrichment score (NES). The entire list of enriched functions is in Table S5. O denotes an old mouse; Y denotes a young mouse; * denotes the mouse used in the analysis. CR: caloric restriction; GH: growth hormone.

To investigate whether transcriptomic changes at the gene-set level are sustained, we compared the GSEA results from samples taken immediately after the attachment period to those after 2 months of detachment. We observed a strong positive correlation between the normalized enrichment scores obtained for the attached and detached groups (Spearman ρ = 0.72, p-value < 10^-16^), and many significantly enriched pathways in the detached analyses were common in identity and direction with those in the attached group, suggesting that transcriptomic changes in liver resulting directly from HPB are largely sustained even after prolonged detachment **(Figure 4B, 6E)**. Of note, the normalized enrichment scores and number of significantly enriched gene-sets were generally decreased in the detached group, despite the same number of samples being analyzed **(Figure 4B)**. This may indicate the gradual reduction of the HPB effect following detachment.

We also conducted global differential gene expression analyses, comparing isochronic and heterochronic samples after the 3-month attachment period or after 2-month detachment. We identified 1044 significantly (adjusted p-value < 0.05) upregulated and 1855 downregulated genes in the post-attachment comparison **(Figure 4C)**, compared to 876 upregulated and 1034 downregulated genes in the detached comparison **(Figure 4D)**. Consistent with the functional enrichment analyses, gene expression changes induced by long-term heterochronic parabiosis right after attachment and after 2-month detachment were positively correlated (Spearman ρ = 0.66, p-value < 1e-308) (**Figure 6D, 7A**). We also observed lower but still significantly positive correlation between short-term and long-term attached samples (Spearman ρ = 0.36, p < 1e-308), as well as between short-term attached and long-term detached samples (Spearman ρ = 0.31, p < 1e-308). This further suggests that HPB induces large-scale, sustained transcriptomic remodeling. Together, these data support that the liver transcriptome is rejuvenated upon HPB, and that this rejuvenation largely persists even after a two-month detachment.

**Figure 7.**
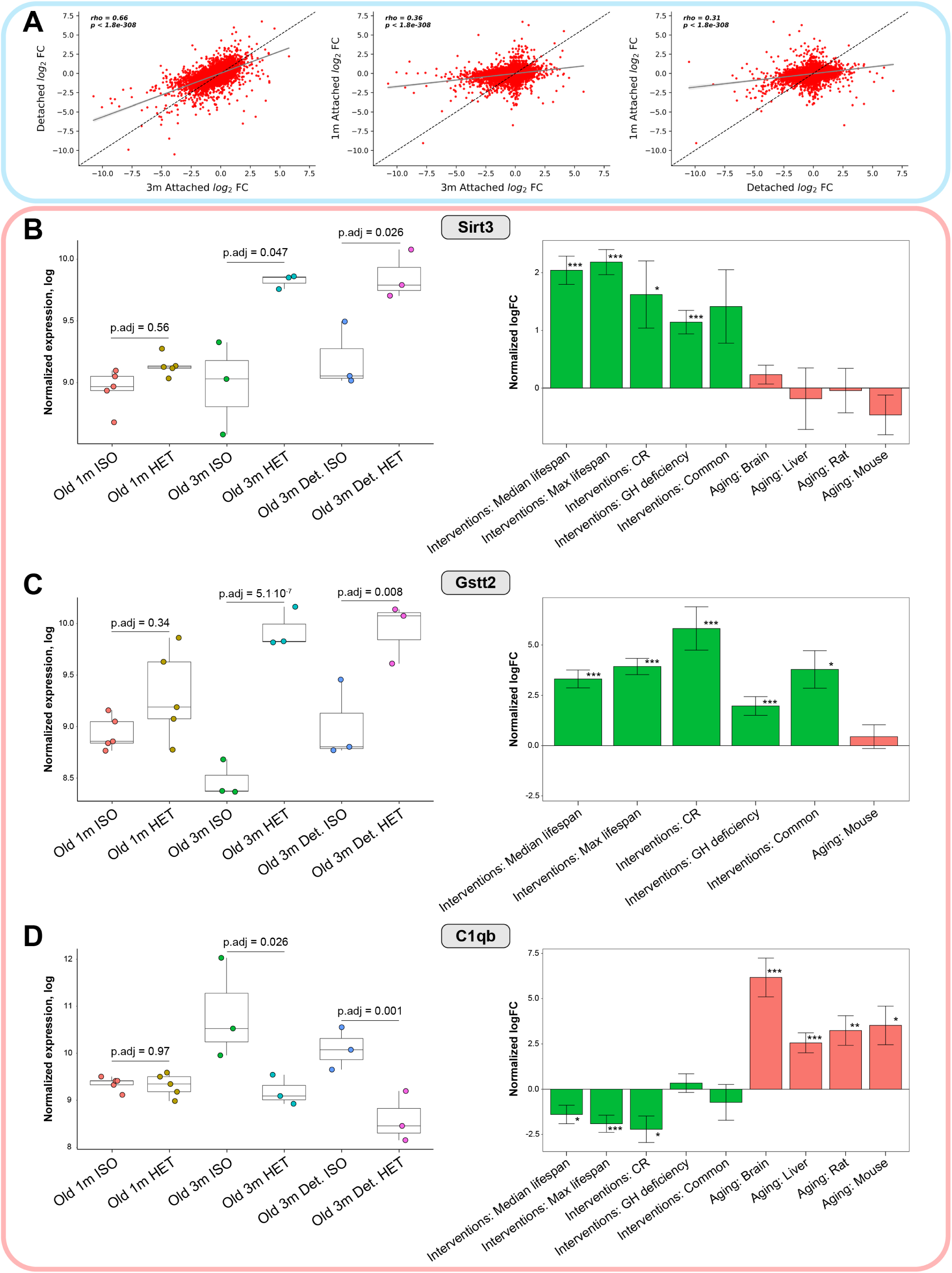
Gene-specific analyses hint at putative rejuvenation mechanisms of long-term HPB. **(A)** Scatterplots depicting the relationship between log_2_ fold-change (log_2_FC) in long-term attached vs. long-term detached (left), long-term attached vs. short-term attached (middle), and long-term detached vs. short-term attached (right). The Spearman correlation coefficient (*rho)* is shown, along with the associated two-tailed p-value (*p)*. The dashed line represents the identity line (same fold change in both conditions). The linear regression line (dark grey) is shown with 95% confidence intervals (light grey). **(B-D)** Expression of *Sirt3* (**B**), *Gstt2* (**C**) and *C1qb* (**D**) in mice subjected to parabiosis (left) and in response to established lifespan-extending interventions and aging (right). For each gene, normalized expression in logarithmic scale is shown across different parabiosis groups: short-term isochronic (Old 1m ISO) and heterochronic mice (Old 1m HET), long-term attached isochronic (Old 3m ISO) and heterochronic mice (Old 3m HET), long-term detached isochronic (Old 3m Det. ISO) and heterochronic mice (Old 3m Det. HET) (left). Adjusted p-values, indicating the difference in expression for each pair of isochronic and heterochronic mice, are included. For every signature associated with lifespan-extending interventions (green) and aging (red), means of normalized logFC or slopes (for signatures of Median and Maximum lifespan) are presented (right). Error bars denote ± 1SE. * p.adjusted < 0.05; ** p.adjusted < 0.01; *** p.adjusted < 0.001.

### Dimensionality reduction reveals intermediate molecular phenotypes resulting from heterochronic parabiosis

Given the high dimensionality of epigenomic and transcriptomic data, we opted to use a well-described dimensionality reduction algorithm, principal component analysis (PCA), to embed and visualize samples in two-dimensional space. When PCA was performed on transcriptomic data from the long-term parabiosis cohort, we observed that heterochronic parabiosis samples clustered between young and old isochronic samples, closer to the young samples than the old **(Figure 5A)**. While clustering of short-term parabiosis also showed an intermediate transcriptomic profile for old heterochronic mice, these samples clustered much closer to the old isochronic group **(Figure 5B)**. These data further suggest that long-term parabiosis is a more potent age-reversal intervention compared to short-term parabiosis, particularly in regard to transcriptomic profiling.

We also conducted PCA analyses on the RRBS methylation data from liver and blood samples **(Figure 5C-H, Figure S5)**. We analyzed methylation at individual CpG sites, as well as through concatenation into two types of genomic regions known to be functionally influenced by cytosine methylation: promoters (**Table S3**) and gene bodies (**Table S4**). Across both tissues, we observed that old heterochronic samples cluster in between old and young isochronic samples when analyzing over 1 million highly-covered CpGs **(Figure 5C-D)**. These results become especially pronounced when collapsing methylation to promoters and gene bodies, where heterochronic samples fall perfectly between the two isochronic groups **(Figure 5E-H)**.

Altogether, dimensionality reduction of epigenomic (RRBS) and RNA-seq data suggests that old heterochronic mice have transcriptomic and epigenetic profiles intermediate between young isochronic and old isochronic mice. This effect is apparent across both the liver and blood, and we moreover observe that long-term parabiosis produces a more profound shift in transcriptomic profiles towards a young state compared to short-term parabiosis. This further indicates that sustained and comprehensive transcriptomic and epigenomic remodeling occurs as a result of long-term, but not short-term HPB.

### Gene expression changes induced by heterochronic parabiosis mirror longevity intervention signatures and oppose aging

To further assess if HPB results in systemic rejuvenation of the transcriptome, we compared the identified gene expression changes with signatures associated with lifespan extension and aging, obtained through a previously published meta-analysis of multiple publicly available datasets **(Figure 6A)**. They include liver-specific gene signatures of individual interventions (such as CR and mutations leading to growth hormone (GH) deficiency), gene expression changes shared across different lifespan-extending interventions, and genes whose expression is associated with the quantitative effect of interventions on maximum and median lifespan. On the other hand, aging signatures reflect age-related transcriptomic changes observed in the liver of mice and rats, as well as multi-tissue age-related alterations in each of these species.

We evaluated the association between parabiosis-induced changes and these signatures using a previously described GSEA-based approach (Tyshkovskiy et al., 2019). In samples taken immediately after detachment or after a prolonged 2-month detachment, we observed a strong and significant negative association with 3 aging signatures, including mouse liver, rat multi-tissue, and mouse multi-tissue aging signatures (adjusted permutation p-value < 5e-4) **(Figure 6B)**. On the other hand, there was a remarkable positive association between HPB in old mice and 4 out of 5 signatures from lifespan-extending interventions, including positive enrichment for median and maximum lifespan signatures and those for caloric restriction and growth hormone deficiency (adjusted permutation p-value < 5e-4). Of note, absolute normalized enrichment scores were consistently decreased in magnitude by 16-24% in post-2- month detached samples compared to samples taken immediately after the long-term parabiosis period **(Figure 6B)**. Consistent with the functional enrichment analysis presented earlier, this suggests that the majority of transcriptomic changes are sustained after the detachment period, but the magnitude of the rejuvenating effect may be gradually diminished over time.

To refine the molecular resolution of longevity effects based on the duration of the parabiosis intervention, we also performed transcriptomic signature analysis for short-term (5-week) parabiosis samples. In accordance with the previous observations, we observed significant positive associations of HPB with several signatures of lifespan extension, while aging signatures demonstrated the opposite effect **(Figure 6C)**. This suggests that short-term parabiosis leads to important transcriptomic changes similar to those produced by established longevity interventions. However, the magnitude of association for short-term parabiosis was reduced by 2-99% compared to long-term attachment. Therefore, long-term parabiosis seems to induce more profound lifespan-extending and rejuvenating effect at the level of gene expression and DNA methylation, highlighting the enhanced effect of long-term HPB on murine lifespan and healthspan as opposed to the short-term treatment.

To investigate mutual associations between the changes induced by heterochronic parabiosis, lifespan extension and aging, we calculated Spearman’s correlation coefficients of the log fold-change of genes for each pair of signatures. Consistent with the GSEA-based association analysis, the effects induced by HPB in old mice clustered together with the changes induced by lifespan-extending interventions and were positively correlated with them (Spearman’s ρ > 0.22; adjusted p-value < 2e-5), but negatively correlated with aging-related changes (Spearman’s ρ < -0.3; adjusted p-value < 3e-9) **(Figure 6D)**. Interestingly, based on our data, the negative association between aging and parabiosis in older animals was even more substantial than between aging and existing lifespan-extending interventions, such as CR (Spearman’s ρ > -0.22).

Finally, to understand which functional gene sets are responsible for the age-reversal and lifespan-extending effects of HPB, we performed functional GSEA for the above-mentioned aging and longevity signatures **(Figure 6E, Table S5)**. We observed that some functions that were upregulated in response to parabiosis, including the TCA cycle, respiratory electron transport and mitochondrial biogenesis, were also upregulated by lifespan-extending interventions, but downregulated during aging. At the same time, functions related to immune response, such as interferon signaling and complement and coagulation cascades, were significantly downregulated both in response to HPB and established lifespan-extending interventions, but upregulated with age. Therefore, HPB in old mice seems to counteract aging by activating genes related to metabolism and cellular respiration while inhibiting inflammatory response, akin to other longevity interventions.

To summarize, we observed that HPB indeed rejuvenated old animals at the systemic level, and this was supported by both epigenetic and transcriptomic remodeling induced by this intervention. Excitingly, signature analyses of HPB profiles appear to recapitulate gene expression effects of established lifespan-extending interventions, pointing to the potential usefulness of this therapy or its derivatives in promoting healthy longevity.

### Individual gene expression dynamics reveal putative longevity-associated mechanisms for HPB

To uncover the potential mechanisms underlying the rejuvenation effects we observed upon long-term HPB, we investigated the individual genes regulated in response to heterochronic parabiosis and their association with longevity and aging based on the gene expression signatures. We found that 27-45% of genes significantly perturbed in response to long-term and short-term HPB were also significantly associated with maximum and median lifespan extension in the same direction (Fisher exact test p-value < 0.03 for all models of HPB). Additionally, there was a significant overlap of genes downregulated in response to long-term heterochronic parabiosis and upregulated with age according to all tested aging signatures (Fisher exact test p-value < 0.004 for long-term attached and detached groups).

Among genes upregulated by long-term HPB in both attached and detached models we identified *Sirt3* (adjusted p-value < 0.048) **(Figure 7B, left)**. *Sirt3* deficiency is known to promote cancer (Gonzalez Herrera et al., 2012) and aging (Benigni et al., 2019; Brown et al., 2013), while its overexpression has resulted in the decrease of reactive oxygen species levels and improvement of regenerative capacity in aged stem cells (Brown et al., 2013). Not surprisingly, its expression in liver is upregulated by many longevity interventions, including CR and growth hormone deficiency (adjusted p-value < 0.049), and is positively associated with the effect of interventions on median and maximum lifespan extension in mice (adjusted p-value < 10^-14^) **(Figure 7B, right)**.

Other important hits, connecting the effects of HPB with mechanisms of established longevity interventions, are genes involved in methionine and glutathione metabolism, such as *Cth* and *Gstt2*, encoding for cystathionine gamma-lyase and glutathione S-transferase theta 2, respectively. Methionine metabolism was shown to play an important role in the regulation of longevity, and the increased blood level of glutathione is known to be an important biomarker of caloric (Lang et al., 1989) and methionine restriction (Richie et al., 1994). Consistent with this data, the expression of *Cth* and *Gstt2* demonstrated significant positive association with the effect of various lifespan-extending interventions (adjusted p-value < 4.10^-12^ for signatures of median and maximum lifespan extension) **(Figure 7C, right)**. At the same time, they were upregulated in response to long-term heterochronic parabiosis, especially in the attached group (adjusted p-value < 0.023) **(Figure 7C, left)**. These findings point to some putative specific longevity mechanisms induced by long-term heterochronic parabiosis.

We also observed a significant increase in *Tert* expression following long-term HPB (adjusted p-value = 4.29.10^-5^), which appeared to be sustained in detached samples (adjusted p-value = 0.0014) **(Figure S6A)**. Although *Tert*, encoding for telomerase reverse transcriptase, doesn’t demonstrate significant upregulation in response to various longevity interventions, the overexpression of this gene by itself has resulted in the lifespan extension in healthy mice without inducing rate of cancer (Bernardes de Jesus et al., 2012).

Among genes downregulated by HPB and longevity interventions but upregulated with age, we identified several involved in the immune response such as *C1qb*, which encodes for one of the complement subcomponents **(Figure 7D)**. This gene appears to be positively associated with aging in liver and brain in both mice and rats (adjusted p-value < 0.05 for all aging signatures). At the same time, its expression in murine liver is downregulated by caloric restriction (adjusted p-value = 0.028) and demonstrates negative association with median and maximum lifespan extension induced by interventions (adjusted p-value < 0.023) **(Figure 7D, right)**. Remarkably, *C1qb* was downregulated in response to long-term heterochronic parabiosis both immediately after attachment and following 2 months of detachment (adjusted p-value < 0.03) **(Figure 7D, left)**, pointing to the potential role of this gene in inducing both the longevity and rejuvenation effects of HPB.

Another interesting hit was *Dnmt3b*, which encodes one of the key enzymes involved in *de novo* methylation of the genome. We observed a significant decrease in *Dnmt3b* expression when comparing isochronic and heterochronic mice from long-term and short-term parabiosis (adjusted p-value < 0.013) **(Figure S6A)**. This effect was also sustained in detached samples (adjusted p-value = 7.3.10^-4^). While *de novo* methylation occurs primarily during embryogenesis and appears to be responsible for a rejuvenation event around the time of gastrulation, this finding may additionally implicate *Dnmt3b* as a key driver in the modulation of epigenetic remodeling induced by HPB (Kerepesi et al., 2021; Okano et al., 1999; Trapp et al., 2021). We also observed significant downregulation of other genes induced by both short-term and long-term HPB, including *Lmna*, *Ly6e* and *Lcn2* **(Figure S6A)**. Remarkably, the expression of these genes was at the same time negatively associated with the effect of longevity interventions on median and maximum lifespan, indicating that they may also mediate the beneficial effect of HPB. Interestingly, for each of these genes, the effect size was strongest in long-term attached samples, intermediate in long-term detached, and lowest in short-term attached samples, again reinforcing the utility of a long-term attachment period for transcriptomic rejuvenation.

Lastly, we also observed strong negative enrichment of senescence-associated secretory phenotype (SASP) genes (**Table S6**) in heterochronic mice immediately after long-term parabiosis (adjusted p-value = 4e-4) (**Figure S7A)**. Expression of SASP-related genes is known to increase with age (Childs et al., 2015), suggesting that HPB ameliorates the pro-inflammatory phenotype of the aging liver. This negative enrichment was also observed in detached heterochronic samples (adjusted p-value = 0.01) and in short-term heterochronic samples (adjusted p-value = 0.008), but the effect in both was noticeably smaller (**Figure S7B-C)**. Gene-specific analyses among several key SASP genes (*Cdkn1a, Cxcl13, Tgfb1*) further revealed significant expression changes between old heterochronic and isochronic mice after 3-month attachment and after 2-month detachment, but not after short-term attachment **(Figure S7D)**.

Together, these results suggest that long-term HPB causes profound transcriptomic change at the level of gene sets and individual genes associated with longevity and rejuvenation. STRING clustering of upregulated and downregulated genes further indicates commonalities in the regulatory gene ensembles that are modified as a result of HPB **(Figure S6B-C)**. The observed shared genes and pathways point to the existence of crucial molecular mechanisms driving the rejuvenation effect of long-term heterochronic parabiosis, some of which have been described in this work.

## DISCUSSION

The history of HPB dates back to the mid-19^th^ century, and it has re-emerged as an important model in aging research since 2005 (Conboy et al., 2005; Conboy and Rando, 2012; Conboy et al., 2013). However, most previous parabiosis studies focused primarily on phenotypes, such as tissue regeneration, brain function and stem cell characteristics. It was enigmatic whether this intervention can systematically reverse the biological age of organisms and, moreover, whether the effects of HPB lasts after a prolonged detachment period. One challenge in this regard has been that the duration of circulatory system attachment in animals was too short to observe concrete and reproducible differences in biological age. By adapting a long-term parabiosis intervention to include detachment and using precise molecular biomarkers of aging and longevity, we demonstrate here for the first time that exposure to the young circulatory system leads to systemic and persistent reversal of biological age, and this effect correlates with a longer lifespan, improved physiological parameters, and a rejuvenated epigenome and transcriptome.

We found that multiple epigenetic clocks point to biological age reversal, and the effect was observed independently based on two different approaches: high-throughput sequencing and microarrays. Importantly, we found that HPB decreases the biological age acceleration (the delta age) even compared to untreated animals, suggesting that this effect is an actual reversal of biological age rather than an amelioration of age acceleration induced by the surgical procedure. It is important to note that while almost all clocks showed significant epigenetic age reversal, some clocks showed only marginal differences. This most often happens when there exists incongruence between the tissues used for clock training and the actual tissues tested, i.e., the blood clock is not as precise in assessing epigenetic age in the liver.

The degree of rejuvenation we observed in our study is consistent with that in previous studies that showed that HPB ameliorates aging phenotypes of tissues other than the blood, such as the muscle, liver and nervous system. This suggests that cells and molecules in the circulatory system contribute to the effect of parabiosis on solid organs and therefore affect cells internal to these tissues (Castellano et al., 2017; Salpeter et al., 2013; Villeda et al., 2011; Villeda et al., 2014). Our finding further fortifies the view that aging is a systemic process (Rando and Wyss-Coray, 2021). Additionally, we observed that gene expression changes induced by HPB in old mice (compared to isochronic controls) were negatively correlated with changes induced by aging, but positively correlated with changes induced by longevity interventions. It should be noted that these effects cannot be explained by the surgical procedure of parabiosis itself, as we subjected all mice to this procedure. Consistent with a recent study examining the effect of short-term parabiosis on single-cell gene expression (Pálovics et al., 2020), our findings indicate that what we observed is in fact a multi-omic, systemic, and sustained biological rejuvenation. Although the effect on the methylation clock showed a decrease of epigenetic age up to 30% by HPB, the lifespan extension effect on the detached mice, while still significant, was not as strong. This is likely due to the fact that mortality in the aged mice was largely affected by tumor incidence, which cannot be changed simply by exposure to the young circulation system (Brayton et al., 2012; Turturro et al., 2002).

Since parabiosis involves direct exchange of young and old blood, it is possible that the presence of some young blood in the liver may contribute to the observed epigenetic age decrease. Based on our results on short-term versus long-term parabiosis, we find that this is unlikely, as the short-term heterochronic old mice should contain similar amounts of chimeric blood as the long-term heterochronic old mice, higher than the amount of the detachment group. However, our results show that the rejuvenation effect of the long-term group is similar to that of the detachment group, while both significantly outperform that of the short-term group. To additionally exclude the possibility of blood contamination as a confounder in our analysis, we calculated the fraction of young blood DNA in the liver of old mice. Previous studies on congenic mouse strains reported that 3-5% of the blood from the parabiont remains in the body after 7 weeks of detachment (Wright et al., 2001). Considering that the residual blood in the liver is roughly 10% of wet weight, and the extracted blood DNA is 2.5-15% of that of the DNA extracted from liver samples of the same weight, the contribution of young blood chimerism is approximately 0.003-0.075% (Davies and Morris, 1993; Satoh, 1979). We cannot preclude a very small number of cells contributing to physiological observations; however, the contribution to DNA yield used for epigenetic analyses should be negligible, since the observed rejuvenation effect is on the order of ∼25%. We also observe intermediate epigenomic and transcriptomic profiles in HPB compared to isochronic controls, highlighting the multi-modal nature of the induced rejuvenation event.

Despite extensive experimentation, the mechanisms of heterochronic parabiosis remain controversial even after decades of research. From our investigation at single-gene resolution, we found that long-term parabiosis causes increased expression of *Sirt3* and genes involved in the methionine and glutathione metabolism, such as *Cth* and *Gstt2*. Remarkably, these genes are also positively associated with the effect of established longevity interventions, pointing to their role in the mediation of healthspan induced by HPB. We also observed decreased expression for genes involved in the immune response, some of which (e.g., *C1qb*) were also downregulated by lifespan-extending interventions but at the same time upregulated with age. Finally, in response to long-term HPB, we detected differential expression of the telomerase reverse transcriptase gene *Tert* and downregulation of the primary *de novo* methylase gene, *Dnmt3b*, which may play a role in the epigenetic remodeling connecting the effect of parabiosis with the observed decrease of DNA methylation age. While it is still difficult to pinpoint the exact causative facets in young blood that provide the rejuvenation effect of HPB, our work provides a platform for deciphering specific gene candidates and pathways associated with this rejuvenation, which may be further investigated in subsequent studies.

It may be that youthful factors in the circulation support a reversal in the biological aging clock. There have been attempts to identify the components involved, and some factors from young blood like oxytocin were reported to recapitulate the physiological rejuvenation effect (Conese et al., 2017; Elabd et al., 2014). It was also reported that, following bone marrow transplantation, the blood of recipients maintains the epigenetic age of the donors, indicating that there may be factors intrinsic to blood cells and their precursors contributing to this effect (Stolzel et al., 2017). Another possibility could be that aged organisms accumulate damage, which can be effectively diluted by young blood during parabiosis experiments. This hypothesis is supported by a recent experiment that involved replacing the blood plasma with saline containing 5% albumin, leading to enhanced muscle repair and hippocampal neuro-regeneration (Mehdipour et al., 2020); however, this has yet to be demonstrated using a longitudinal model. It is critical to note that there has been no evidence from aging clocks favoring either of these two hypotheses. Therefore, it would be interesting to use molecular aging biomarkers to clarify the exact causes of this rejuvenation effect.

Overall, our findings reveal a robust decrease in epigenetic age when aged mice are attached to the young, which remains strongly significant after detachment. This is coupled with a strong positive correlation between parabiosis and longevity interventions and a negative correlation between parabiosis and aging based on gene expression signatures. Gene set and pathway analyses revealed a positive enrichment of TCA cycle, oxidative phosphorylation, mitochondrial biogenesis and fatty-acid metabolism pathways as well as negative enrichment of interferon-gamma and inflammatory response pathways in heterochronic mice. Additionally, several longevity-mediated genes were identified, including *Gstt2*, *Sirt3*, *C1qb* and *Tert*, that showed sustained expression changes following long-term parabiosis. Importantly, we validate here that long-term (3-month) parabiosis is much more effective at rejuvenating the transcriptome and epigenome of mice compared to short-term (5-week) parabiosis. This long-lasting rejuvenation effect of HBP in old mice leads to the extension of healthspan and lifespan even after this traumatic surgery, and produces molecular profiles intermediate between young and old mice. At the moment, several interventions or biological events have been shown to reverse the epigenetic age, including rejuvenation during early embryogenesis, rejuvenation by pharmaceutical treatments and rejuvenation via reprogramming factor expression. Given this, it would be interesting to investigate molecular commonalities resulting from these interventions, helping to design perturbations that recapitulate the rejuvenation effect without requiring a complex and impractical surgery such as parabiosis. To conclude, our results indicate that biological age and molecular damage can be systemically reversed in a sustained manner following exposure to young circulation, and open exciting new avenues for research on parabiosis and its derivatives for organismal rejuvenation.

## MATERIALS AND METHODS

### Mouse models

Female C57BL/6J (Wild-type, Wt) mice were used at 3 months and 20 months of age. The selection of female mice was based on better temperament when conjoined and survival rates in females compared to males after the surgeries. All animal care followed the guidelines and was approved by the Institutional Animal Care and Use Committees (IACUCs) at Duke Medical Center. All old mice (20m) were acquired from the NIA aging colony.

### Parabiosis/detachment surgery

Parabiosis methods were performed as previously described (Baht et al., 2020), with modifications to the length of the protocol. In brief, parabiosis surgery was carried out using 3 and/or 20-month-old female mice. Mouse pairs were selected at random following a prescreening to minimize differences in body size. All pairs were maintained together for 3 months, then euthanized or detached for further analysis. The detachment surgery was performed under isoflurane anesthesia, separating the pairs by reversing the attachment site. Once both mice were separated, skin and fascia were sutured closed. For longevity analysis, all detached mice were allowed one month of recovery before entry into the study in an attempt to account for any premature deaths due to surgical complications. Experimental groups for the longevity study were isochronic old (Old ISO or O:O) and heterochronic old (Old HET or Y:O). Mice used for longevity studies were not used for healthspan studies to prevent confounding effects, and allowed to live until natural death or recommendation of euthanasia by a veterinarian. For healthspan studies, all mice were detached after the 3-month parabiosis period and phenotypic and functional data collection started one month after detachment to allow for surgical recovery. Experimental groups included isochronic young (Young ISO or Y:Y), isochronic old (Old ISO or O:O) and heterochronic old (Old HET or Y:O). The same groups were used for detachment cohorts. For epigenetic analyses, all mice were anastomosed for 3-months then analyzed while still attached or analyzed two months after detachment. Experimental groups included isochronic young (Young ISO or Y:Y), isochronic old (Old ISO or O:O) and heterochronic old (Old HET or Y:O). The same groups were used for detachment cohorts and annotated with a “D” or “Det.” on our data sets and figures. We have previously shown a shared blood supply at 5 weeks (Vi et al., 2018) and now show continued sharing at 3 months after surgery using flow cytometry as shown in **Figure S2**.

### Genomic DNA and total RNA isolation

Five DNA samples were taken from each group for microarray analysis, 5-7 samples per group for RRBS, and 3 samples for RNA-seq analysis described in **Table S1**. DNA from samples was isolated by using the DNeasy Blood & Tissue Kit (Qiagen 69506), and then eluted from columns in 100 µl of 10 mM Tris-HCl buffer, pH 8.0. 2 µl of RNase A (Life Technologies) was added to each sample. Samples were incubated at room temperature for 2 min, and isolated genomic DNA was purified by using the Genomic DNA Clean & Concentrator™-10 kit (Zymo D4011). DNA was eluted in 25 µl of 10 mM Tris-HCl buffer, pH 8.0 and quantified using a Qubit 2.0 (Life Technologies AM2271). RNA from liver samples RNA was eluted by the Invitrogen Ambion RNAqueous Total RNA Isolation Kit (Invitrogen AM1912) in 60 µl of nuclease free water and quantified using Qubit 2.0 (Life Technologies).

### RRBS library preparation and epigenomic data processing

RRBS libraries were prepared following the protocol previously established (Petkovich et al., 2017), using 100 ng of DNA for each sample. Each library included 10 samples. To avoid an overlap of batch effect and age-related changes, we randomized samples into 9 separate libraries (**Table S1**). Liver and blood samples were analyzed separately. The libraries were sequenced using an Illumina HiSeq2500, with 150bp paired-end reads. 20% of mouse genomic DNA library was spiked in to compensate for low complexity of the libraries. RNA samples were sequenced separately. Adapter removal and quality trimming for the analysis of DNA methylation reads were performed using TrimGalore v0.4.1 following previously established protocols (Krueger and Andrews, 2011; Meer et al., 2018; Petkovich et al., 2017). For the genomic DNA methylation analyses, the trimmed reads were mapped to the mouse genome (GRCm38.p2/mm10) using Bismark v0.15.0. (Krueger and Andrews, 2011) Coverage files outputted by Bismark were then used for further analyses. To ensure high accuracy and precision in methylation levels, CpG sites that had less than 10-read-coverage were filtered out. When combining liver and blood samples into one dataframe, a total of 1,014,243 CpGs were covered at a depth greater than 10x across all samples. For promoter methylation analyses, the promoter region of each gene was determined as [-1500, +500] bp from the transcription starting site following the direction of transcription by using the RefSeq annotation file MmRefSeqTSS.sga, found at https://ccg.epfl.ch/mga/mm10/refseq/refseq.html. Mean methylation level of CpG sites in the promoter regions was calculated for each gene by taking the average of all CpGs covered in a particular promoter. To reduce noise from low coverage, only promoters that had at least 5 CpG sites with valid data were retained. This resulted in coverage of 11,842 promoters across all liver and blood RRBS samples. Gene body methylation analyses were conducted by concatenating all CpGs within a particular gene body (from transcription start site to transcription end site) and averaging methylation, just as in promoter methylation analyses (again with a 5 CpG filter). This resulted in coverage of 13,811 gene bodies across all blood and liver samples.

### Application of mouse epigenetic clocks to RRBS datasets

Four clocks were applied to the RRBS dataset to characterize epigenetic age, including two whole-lifespan multi-tissue clocks, a blood-based clock (Petkovich et al., 2017), and a recently developed single-cell clock (Trapp et al., 2021), modified to accommodate bulk data **(Figure 2)**. Epigenetic clock analyses for conventional elastic-net based clocks (Meer et al., 2018; Petkovich et al., 2017; Thompson et al., 2018) were performed exactly following the descriptions in the original research articles.

In the case of the single-cell clock (*scAge)*, several modifications were applied compared to its original application (Trapp et al., 2021). Indeed, in the original *scAge* application to sparse single-cell data, single-cell profiles were predominantly binary, but this is not the case with bulk RRBS data. Bulk methylation levels are often fractional, given that several dozen reads coming from different cells usually cover a single CpG. Therefore, to account for this biologically meaningful fractional methylation distribution, no forced binarization of RRBS profiles was performed prior to running the algorithm. Additionally, the core algorithm was modified to measure probabilities based on the absolute distance between observed and predicted methylation levels for any given age, without an inherent requirement for binary observed methylation values. Moreover, this enabled epigenetic age predictions without modification to the input data. Given the targeted nature of RRBS, the top 25% age-associated CpGs per sample were used for predictions. As training data to the modified *scAge* framework, we utilized bulk C57BL/6J RRBS data from the Thompson et al. 2018 study. Two CpG-specific reference tables were created: (1) bulk blood RRBS methylation profiles from 50 normally-fed C57BL/6J mice across 1,202,751 CpG sites and (2) bulk liver RRBS methylation profiles from 29 normally-fed C57BL/6J mice across 1,042,996 CpG sites on the positive strand. Based on dimensionality reduction analyses, two blood samples and one liver sample that were evident outliers were removed from downstream epigenetic aging analyses **(Figure 5)**. Welch’s one-tailed t-test assuming unequal variances was used for statistical testing.

### Application of mouse epigenetic clocks to the DNA microarray dataset

We analyzed 5 samples in each group with the recently developed Infinium array HorvathMammalMethylChip40 (Lu et al., 2021). Four clocks were applied in our analysis, including 2 clocks based on the mouse liver (Mouse Liver and Mouse Liver Development), a universal mammalian clock based on age relative to maximum lifespan, and a universal mammalian clock based on log-linear transformed age **(Figure 3)**. The Mouse (Liver Development) clock was similar to the Mouse (Liver) clock, except that sites were sub-selected based on changes occurring during development in this organ. The epigenetic age data are in **Table S2**. Welch’s one-tailed t-tests assuming unequal variances were used for statistical testing.

### RNA-seq analysis of heterochronic parabiosis

Paired-end RNA sequencing for mouse liver samples was performed on an Illumina NovaSeq 6000 S4 platform with 100 bp read length. Reads were mapped with STAR (version 2.5.2b) (Dobin et al., 2013) and counted via featureCounts (version 1.5) (Liao et al., 2014). Data was passed through RLD transformation (Love et al., 2014), and genes with |transformed normalized count| < 1 were filtered out. This left 22,772 genes, which were used for dimensionality reduction and further analyses **(Figures 4-7)**. For signature association analysis and identification of longevity-associated genes perturbed by heterochronic parabiosis, we filtered out genes with low number of reads, keeping only the genes with at least 10 reads in at least 50% of the samples, which resulted in 12,374 detected genes according to Entrez annotation.

### Differential gene expression analyses

Differential expression analyses were conducted using DEseq2 on normalized count data. All samples were concatenated to a single data frame and normalized jointly prior to analysis, in order to minimize batch-effect bias. 15,481 valid genes were obtained when conducting differential expression analysis on long-term parabiosis samples comparing old isochronic and old heterochronic mice. Similarly, 14,522 valid genes were obtained when conducting DEA on long-term detached parabiosis samples comparing old detached isochronic and old detached heterochronic mice. These data were used to generate volcano plots for attached and detached samples in **Figure 4**. Volcano plots were generated using the package *bioinfokit* 2.0.6.

For signature association analyses **(Figure 6)** and investigation of longevity-associated genes regulated by HPB **(Figure 7)**, differential expression of genes in response to heterochronic parabiosis compared to isochronic parabiosis was further analysed using edgeR (Robinson et al., 2010) separately for long-term attached, long-term detached and short-term attached models. Obtained p-values were adjusted for multiple comparison with Benjamini-Hochberg method (Benjamini and Hochberg, 1995). The RLE method was used to obtain normalized expression of genes (Anders and Huber, 2010).

### Dimensionality reduction

To identify trajectories of molecular profiles resulting from HPB, we conducted dimensionality reduction via Principal Component Analyses (PCA) on gene expression (RNA-seq) and methylation (RRBS) data **(Figure 5)**. For gene expression analyses, we used normalized, RLD-transformed gene count values from the liver across 22,272 genes in both short-term and long-term parabiosis samples **(Figure 5A-B)**. For sequencing-based methylation data, we used information at 1,014,243 highly-covered (>10x) CpG sites in liver and blood samples. This was then collapsed into 11,842 promoters and 13,811 gene bodies, each covered by at least 5 CpGs to ensure reliable mean methylation measurements. The percentage of the variance explained by the first two principal components was extracted and is shown across **Figure 5**.

### Association of signature-based gene expression changes induced by parabiosis, aging and lifespan extending interventions

Gene expression signatures of lifespan-extending interventions, including: signatures of caloric restriction (CR), signatures of growth hormone deficiency, common gene expression changes across different interventions, and signatures associated with the effect of interventions on median and maximum lifespan, were obtained from (Tyshkovskiy et al., 2019). Aging signatures, including the liver signature as well as the mouse and rat multi-tissue signatures, were obtained via a meta-analysis of age-related gene expression changes. Association of gene expression log fold-changes associated with parabiosis, lifespan extension and aging were assessed using correlation and GSEA (Subramanian et al., 2005). For the correlation analysis, a Spearman correlation metric was calculated using the top 200 statistically significant genes for each pair of signatures. Clustering was subsequently performed with a complete hierarchical approach **(Figure 6)**.

For the GSEA-based association analysis, we utilized an algorithm developed in (Tyshkovskiy et al., 2019). First, for every signature we specified 250 genes with the lowest p-values and divided them into up- and downregulated genes. These lists were subsequently considered as gene sets. Then, we ranked genes differentially expressed in response to parabiosis based on their p-values, calculated as described in the functional enrichment methods section. Afterwards, we calculated normalized enrichment scores (NES) separately for up- and downregulated lists of gene sets as described in (Tyshkovskiy et al., 2019) and defined the final NES as a mean of the two. To calculate statistical significance of obtained NES, we performed permutation testing where we randomly assigned genes to the lists of gene sets, maintaining their size. To get the p-value of the association between parabiosis and a certain signature, we used 5,000 permutations and calculated the frequency of random final NES that are larger in magnitude than the observed final NES. To adjust for multiple testing, we performed a Benjamini-Hochberg correction (Benjamini and Hochberg, 1995). Final NES for association of HPB response with various longevity and aging signatures were used to generate barplots **in Figure 6**.

Overlap between significant differentially expressed genes in response to HPB and lifespan-extending interventions or aging was assessed using Fisher’s exact test separately for sets of up- and downregulated genes.

### Functional enrichment analysis

For the identification of enriched functions distinguishing isochronic and heterochronic mice, we performed functional GSEA (Subramanian et al., 2005) on a pre-ranked list of genes based on log_10_(p-value) corrected by the sign of regulation, calculated as:

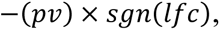

where *pv* and *lfc* are p-value and logFC of a certain gene, respectively, obtained from *edgeR* output, and *sgn* is the signum function (equal to 1, -1 and 0 if value is positive, negative or equal to 0, respectively). REACTOME, KEGG and GO BP ontologies from the Molecular Signature Database (MSigDB) were used as gene sets for GSEA. The GSEA algorithm was performed separately for long-term attached and detached models via the *fgsea* package in R with 5000 permutations. A q-value cutoff of 0.1 was used to select statistically significant functions.

A similar analysis was performed for gene expression signatures associated with aging and lifespan extension. A heatmap colored by NES was built for manually chosen statistically significant functions **(Figure 6)**. Clustering of functions was performed with hierarchical complete approach and Spearman correlation distance.

For parabiosis samples, we also conducted additional GSEA using the Hallmark gene sets, downloaded from MSigDB. We used a Python GSEA implementation, *gseapy* 0.10.5, to perform gene set enrichment. Only Hallmark gene sets with Benjamini-Hochberg adjusted p-values below 0.05 are shown in **Figure 4**. GSEA was conducted separately for long-term attached and long-term detached samples, with 5000 gene set permutations for statistical testing. Hallmark terms significantly enriched in the same direction across both attached and detached comparisons are highlighted in red.

### Cage activity

Voluntary activity was tested in the open field arena (Omnitech, Columbus, OH) illuminated at 340 lux as previously described by Huffman et al. (Huffman et al., 2021). Mice were placed individually into the open field and baseline locomotion was monitored over 60 min.

### Statistical and computational analyses

Computational analyses were performed in Python 3.9.2, running with *numpy* 1.20.0 and *pandas* 1.2.4. Statistical analyses for epigenetic ages were performed with one-tailed Welch’s t-test, implemented in *scipy* 1.32.0. Statistical analyses for gene expression were performed with custom models from *DEseq2* 3.13 and *edgeR* 3.34.1 in R 4.0.3 (Robinson et al., 2010). Statistical analyses of gene expression signatures, gene set enrichment analyses, and epigenetic age profiling were performed as documented above.

For the statistical evaluation of body composition, activity, and food consumption data (**Figure 1c-f**, **Figure S3a-b**) we used a custom test based on random simulation. We generated a randomly simulated point for each time point of old isochronic samples using the average and standard deviation of the old isochronic samples of each time point. In a similar way, we generated another randomly simulated point for each time point of old heterochronic samples. This process resulted in a simulated time series for old heterochronic samples and simulated time series for old isochronic samples. We assessed these time series by summarizing differences for simulated old isochronic - old heterochronic pairs for all time points. Repeating the whole procedure 100,000 times, we calculated the p-value as the proportion of the cases when the difference between both time series was > 0, if the null hypothesis was ‘old isochronic > old heterochronic’; and < 0, if the null hypothesis was ‘old isochronic < old heterochronic’. If the p-value was less than 0.05, we rejected the null hypothesis and considered the difference significant. The null hypothesis was ‘old isochronic > old heterochronic’ in the case of body weight, lean mass, distance and vertical activity, and ‘old isochronic < old heterochronic’ in the case of fat mass and food consumption.

### Data Availability

The data obtained in this study will be deposited to GEO with the accession number GSEXXXXXX.

## Supporting information

Supplementary Images

## Acknowledgements

J.P.W. was supported by NIH grants K01AG056664 and R21AG065943. V.N.G. was supported by NIH grants R01AG067782, P01AG047200 and R01AG065403. D.E.L was supported by NIH training grant T32HL007057. S.H. acknowledges support from the Milky Way Research Foundation and the Epigenetic Clock Development Foundation, and A.Tyshkovskiy and S.E.D. acknowledges support from the Interdisciplinary Scientific and Educational School of Moscow University “Molecular Technologies of the Living Systems and Synthetic Biology”. We thank the Duke Behavioral Core for support on this project. We also thank Tiamat Fox for help with schematic figures. Figure icons were partially created with BioRender.com.

## Author contributions

B.Z., V.N.G. and J.P.W. conceived the project. D.E.L. and J.P.W. carried out animal experiments. A.Trapp and B.Z. designed figures. A.Trapp, A. Tyshkovskiy and B.Z. performed gene expression analyses. B.Z., A.Trapp, and C.K. conducted DNA methylation clock analyses based on RRBS, and B.Z., A.T.L. & S.H. based on microarrays. A.V.S contributed to enrichment analyses. S.E.D. contributed to data interpretation. V.N.G. and J.P.W. supervised the project. B.Z., A.Trapp, V.N.G. and J.P.W. wrote the manuscript with input from all co-authors.

## Competing interests

The authors do not report any conflicts of interest.

## SUPPLEMENTARY FIGURE LEGENDS

**Figure S1. Timeline of molecular and physiological profiling.**

**Related to Figure 1**

**(A)** Schematic timeline of the molecular (transcriptomic & epigenetic) and physiological profiling presented in this study. Samples were taken immediately after the parabiosis period (month 3), or after a 2-month detachment period (month 5). Physiological profiling was performed before the parabiosis experiment (month 0), and then every month from month 4 to month 10.

**(B)** Changes in food consumption resulting from parabiosis. Lines depict mean changes with 95% confidence intervals. Individual observations are shown as points. Young isochronic (Young ISO) mice are shown in blue, old heterochronic (Old HET) mice are shown in purple, and old isochronic (Old ISO) mice are shown in red. Dashed line depicts time of detachment.

**(C)** Changes in vertical activity resulting from parabiosis. Lines depict mean changes with 95% confidence intervals. Individual observations are shown as points. Legend is the same as (B).

**Figure S2. Blood mixture analyses after 3 months of parabiosis.**

**Related to Figure 2**

**(A)** Blood sharing in old mice based on 2 flow cytometry analyses of whole blood gated on GFP (+), (x-axis) and CD45 (+) (y-axis) cells.

**(B)** Histogram of GFP counts across intensity. Blood was sampled from old wild-type mice anastomosed to young GFP mice during the final week of the 3-month protocol to ensure blood sharing in our long-term HPB. Pie plot represents percent of GFP (+) and (-) cells.

**Figure S3. Relationship of DNAm age across tissues and platforms.**

**Related to Figure 2, 3**

**(A)** Scatterplots highlighting the association of liver and blood DNAm predictions based on RRBS sequencing across 6 different group: young isochronic (Young ISO), young isochronic detached (Young ISO Det.), old heterochronic (Old HET), old isochronic (Old ISO), old heterochronic detached (Old HET Det.), and old isochronic detached (Old ISO Det.). Each point depicts the mean DNAm prediction for a particular tissue for that group. The clock used for predictions is shown in the top part of each panel. The Pearson correlation *(r)* is shown in each panel, along with its associated two-tailed p-value. Linear regression lines (dark grey) with 95% confidence intervals (light grey) are shown.

**(B)** Scatterplots highlighting the association of liver DNAm predictions based on RRBS sequencing and methylation array profiling across 6 different groups (same legend as (A)) Each coordinate for a point depicts the average DNAm prediction for that group in a particular platform. The clock used for RRBS predictions is shown in the top part of each panel. The mouse liver clock was used for array predictions. The Pearson correlation *(r)* is shown in each panel, along with its associated two-tailed p-value. Linear regression lines (dark grey) with 95% confidence intervals (light grey) are shown.

**Figure S4. Mean methylation in array and RRBS methylation profiles. Related to Figure 2, 3.**

**(A)** Mean liver methylation assayed by the Mammalian Methylation Array (HorvathMammalMethylChip40), both based on all sites in the array (left, Global methylation) and only sites in CpG islands (right, CpG island methylation). *n* = 5 samples per group.

**(B)** Mean global methylation (top), promoter methylation (middle), and gene body methylation (bottom) of liver (left) and blood (right) RRBS samples. Mean methylation was assayed across 1,014,243 CpGs (top), 11,842 promoters (middle), and 13.811 gene bodies (bottom). *n* = 4-7 samples per group.

Two-tailed Welch’s t-test was used for statistical analysis. Schematic tissues in each panel indicate the source of the sample.

**Figure S5. Dimensionality reduction highlights tissue specificity of RRBS-based epigenomic profiles.**

**Related to Figure 5.**

**(A)** Principal component analysis (PCA) of CpG methylation across 1,014,243 CpG sites in *n* = 36 liver samples and *n* = 32 blood samples, with 6 different groups in both tissues: young isochronic (Young ISO), young isochronic detached (Young ISO Det.), old heterochronic (Old HET), old isochronic (Old ISO), old heterochronic detached (Old HET Det.), and old isochronic detached (Old ISO Det.).

**(B)** PCA of CpG methylation across 11,842 promoters in *n* = 36 liver samples and *n* = 32 blood samples. Promoter methylation was calculated by averaging methylation across CpGs within [-1,500 bp, 500 bp] of the transcription start site (TSS) of a gene, with only promoters containing >5 CpGs considered for downstream analysis. The legend described in (A) applies also to this panel.

**(C)** PCA of CpG methylation across 13,811 gene bodies in *n* = 36 liver samples and *n* = 32 blood samples. Gene body methylation was calculated by averaging methylation across CpGs located between the TSS and transcription end site (TES) of a gene, with only genes containing >5 CpGs considered for downstream analysis. The legend described in (A) applies also to this panel. In all these PCA plots, tissue of origin is clearly the greatest source of variation (44-91%).

**Figure S6. Transcriptomic changes resulting from HPB.**

**Related to Figure 7**

**(A)** Boxplots of RLD-transformed, log-normalized count of several genes (top boxes) across 6 groups (from left to right: old short-term isochronic, old short-term heterochronic, old long-term isochronic, old long-term heterochronic, old long-term isochronic detached, and old long-term heterochronic detached). The log_2_ fold-change (log_2_FC) and associated p-value are shown for each comparison.

**(B)** STRING network representation of significantly downregulated genes in livers of old heterochronic mice (*n* = 421 genes) compared to isochronic mice immediately after detachment or after a 2-month detachment period (protein-protein interaction q-value < 1e-16). Genes were filtered based on directionality and absolute value of the log_2_FC.

**(C)** STRING network representation of significantly downregulated genes in livers of old heterochronic mice (*n* = 337 genes) compared to isochronic mice immediately after detachment or after a 2-month detachment period (protein-protein interaction q-value < 1e-16). Genes were filtered based on directionality and absolute value of the log_2_FC.

**Figure S7. SASP enrichment and differential expression across long-term and short-term HPB.**

**Related to Figure 7**

**(A-C)** Running enrichment score for the senescence-associated secretory phenotype (SASP) gene set, comparing in **(A)** old isochronic and old heterochronic mice from the long-term HPB, in **(B)** old detached isochronic and old detached heterochronic mice from long-term HPB, and in **(C)** old detached isochronic and old detached heterochronic mice from short-term HPB. The p-values for the gene set enrichment, along with the adjusted p-value are shown in each panel. Positions of individual genes in the gene set are shown as black bars near the bottom of each panel. Long-term attached HPB shows the greatest negative enrichment for heterochronic samples, followed by detached long-term HPB and attached short-term HPB, respectively.

**(D)** Boxplots of RLD-transformed, log-normalized count of five SASP genes across 6 groups (from left to right: old short-term isochronic, old short-term heterochronic, old long-term isochronic, old long-term heterochronic, old long-term isochronic detached, and old long-term heterochronic detached). The log_2_ fold-change (log_2_FC) and associated p-value are shown for each comparison.

## Notes

### Competing Interest Statement

The authors have declared no competing interest.

