## Supplementary figures and images for "Multi-omic rejuvenation and lifespan extension upon exposure to youthful circulation"

### Supplementary Images

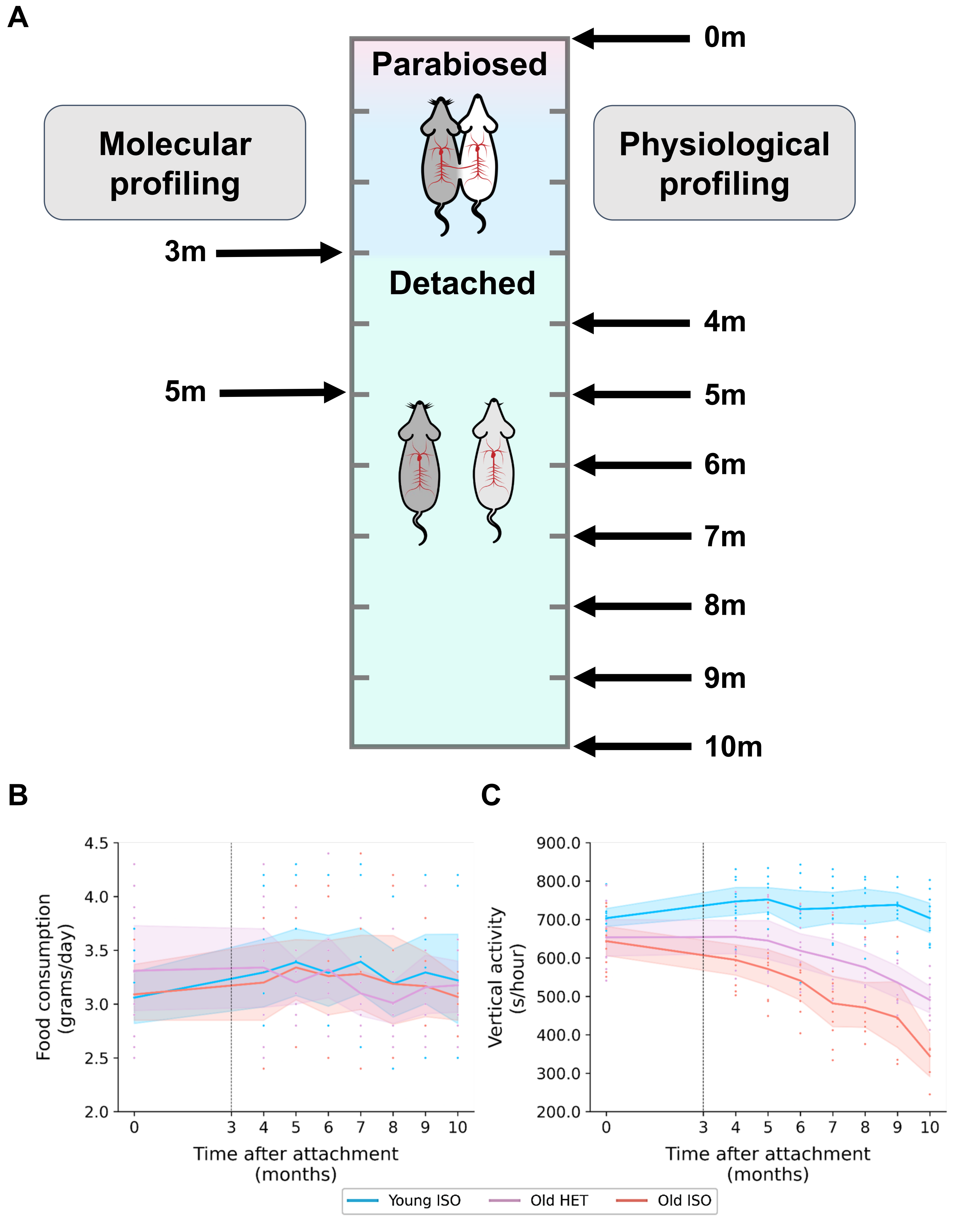

#### Figure S2

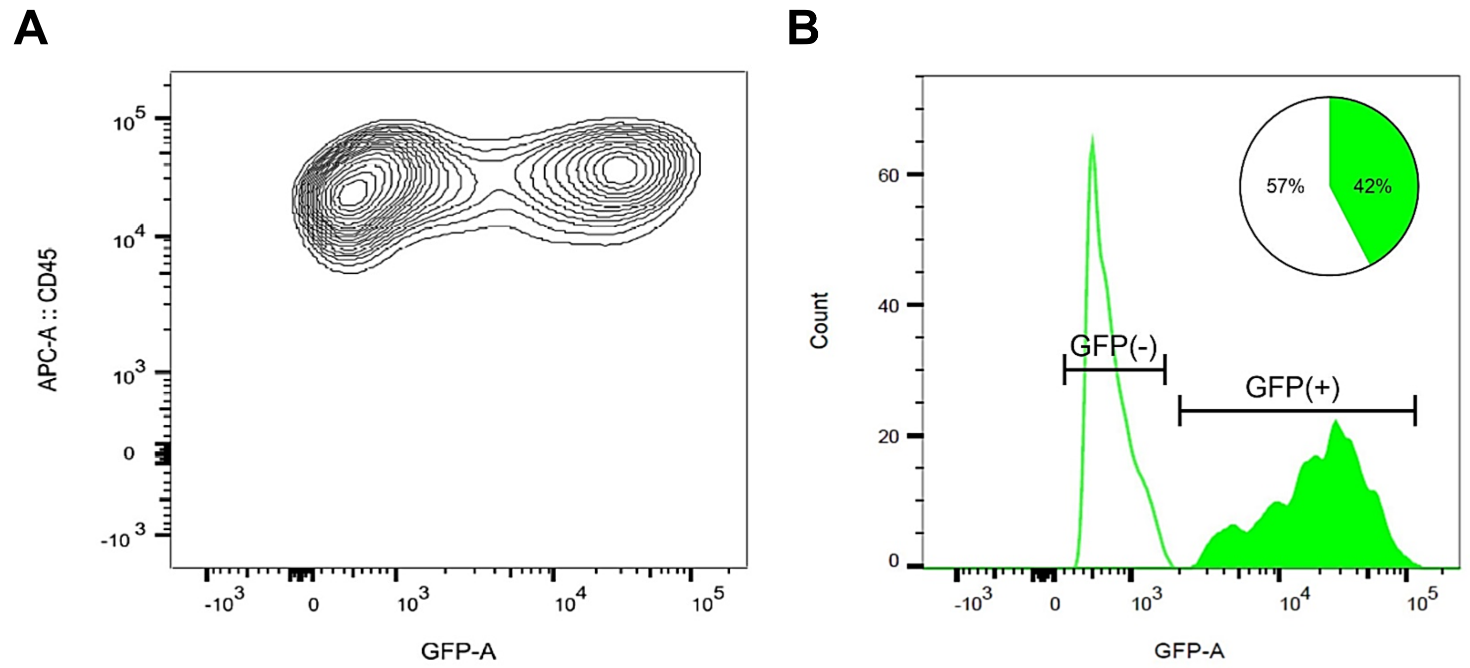

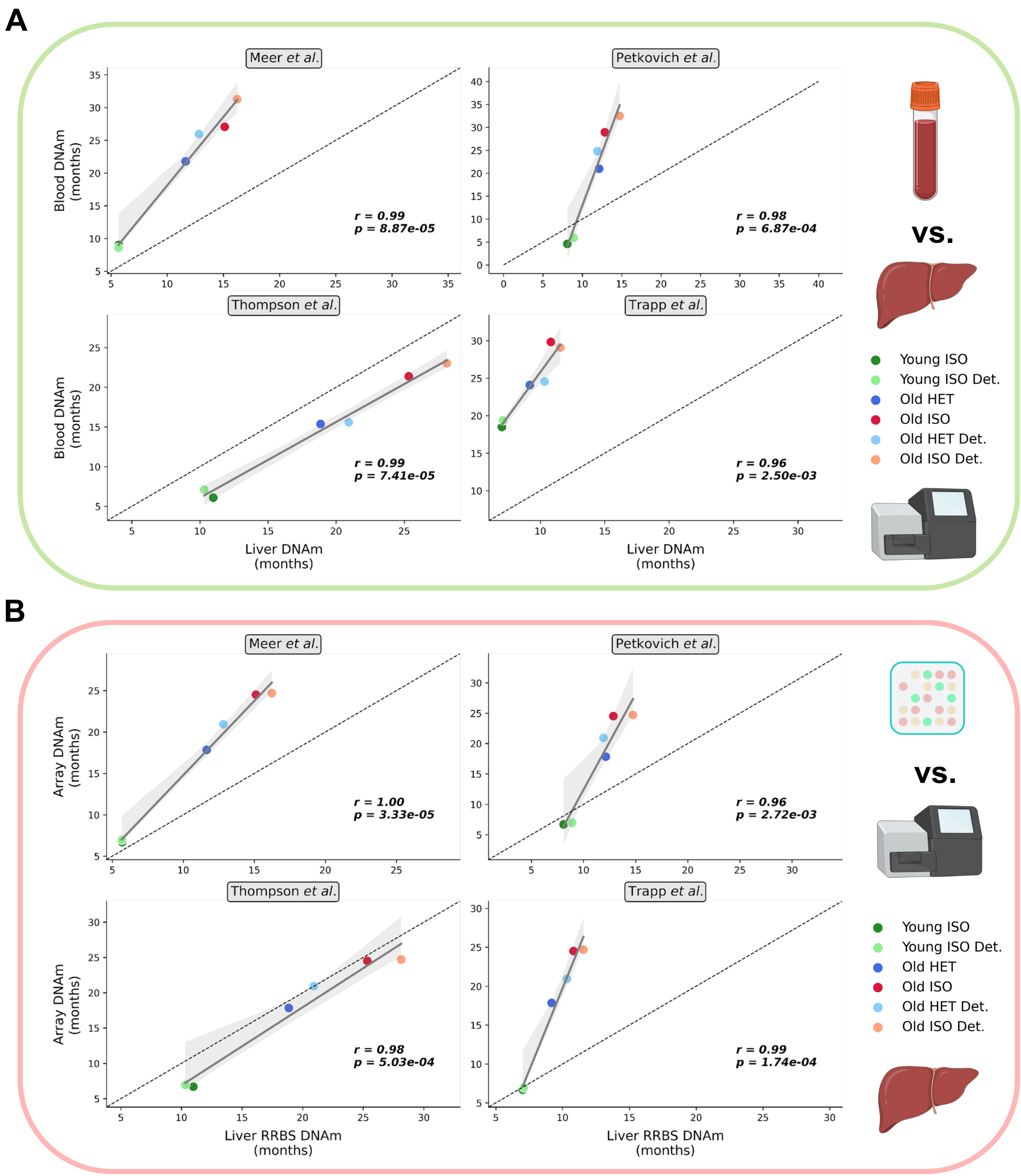

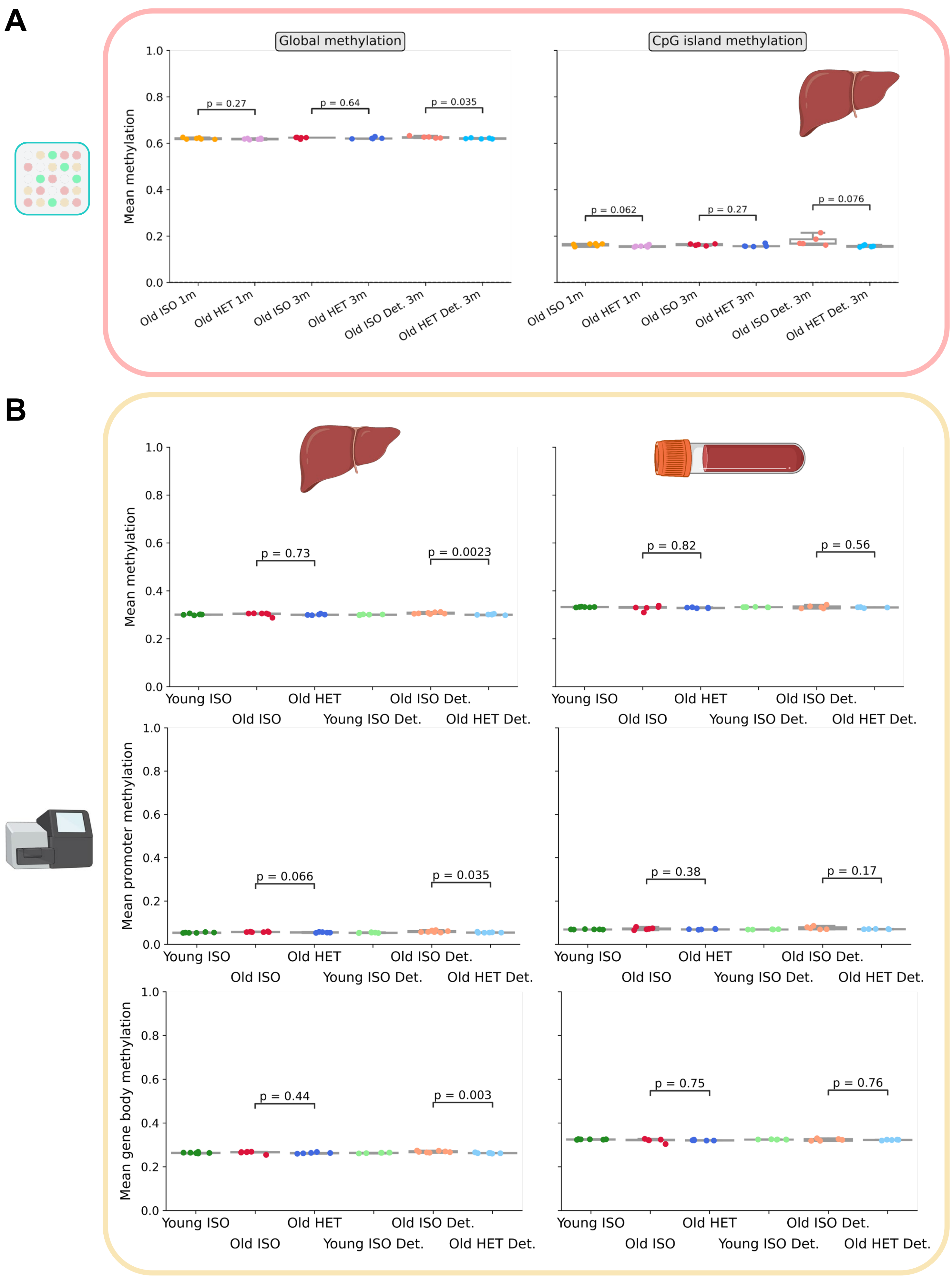

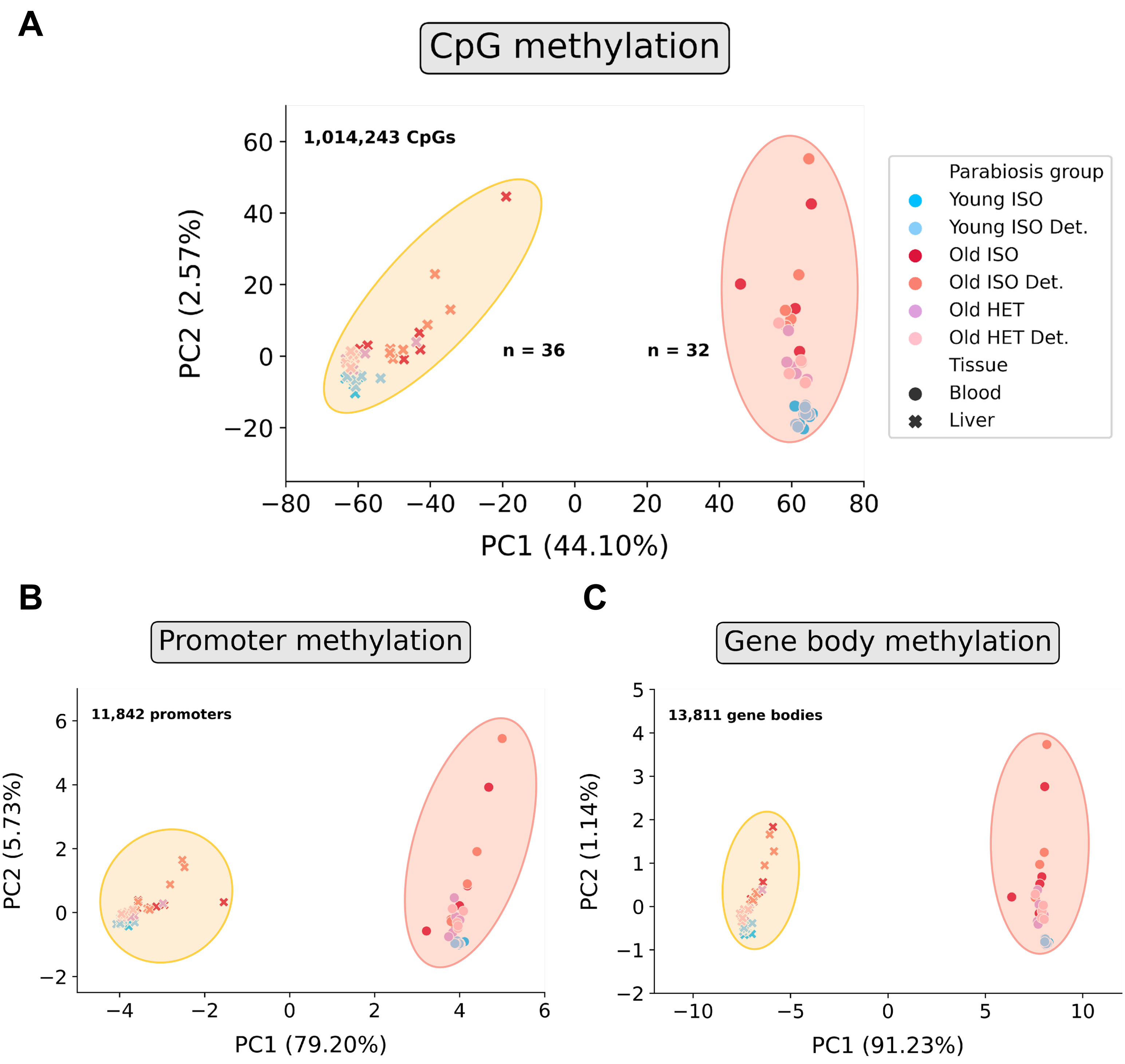

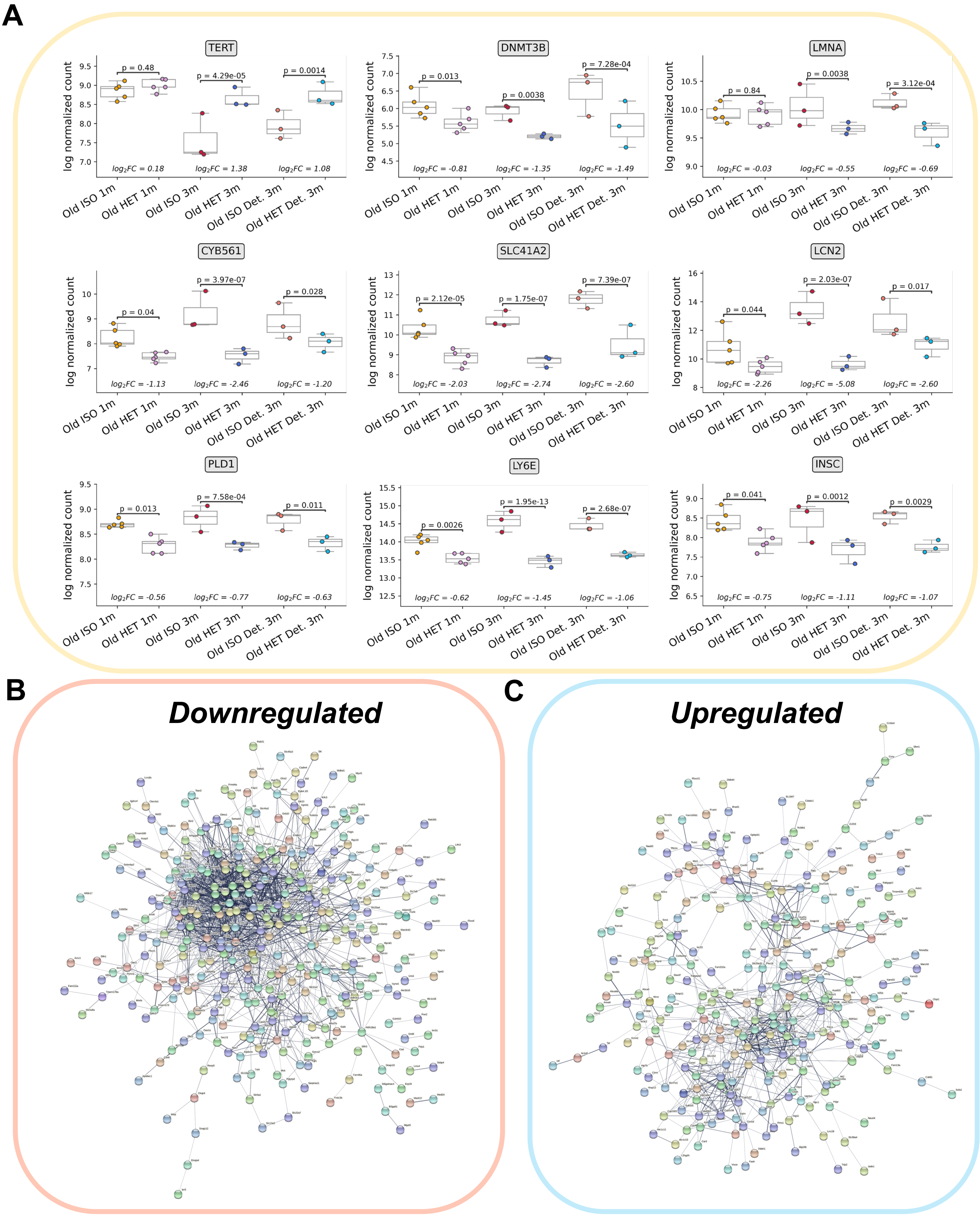

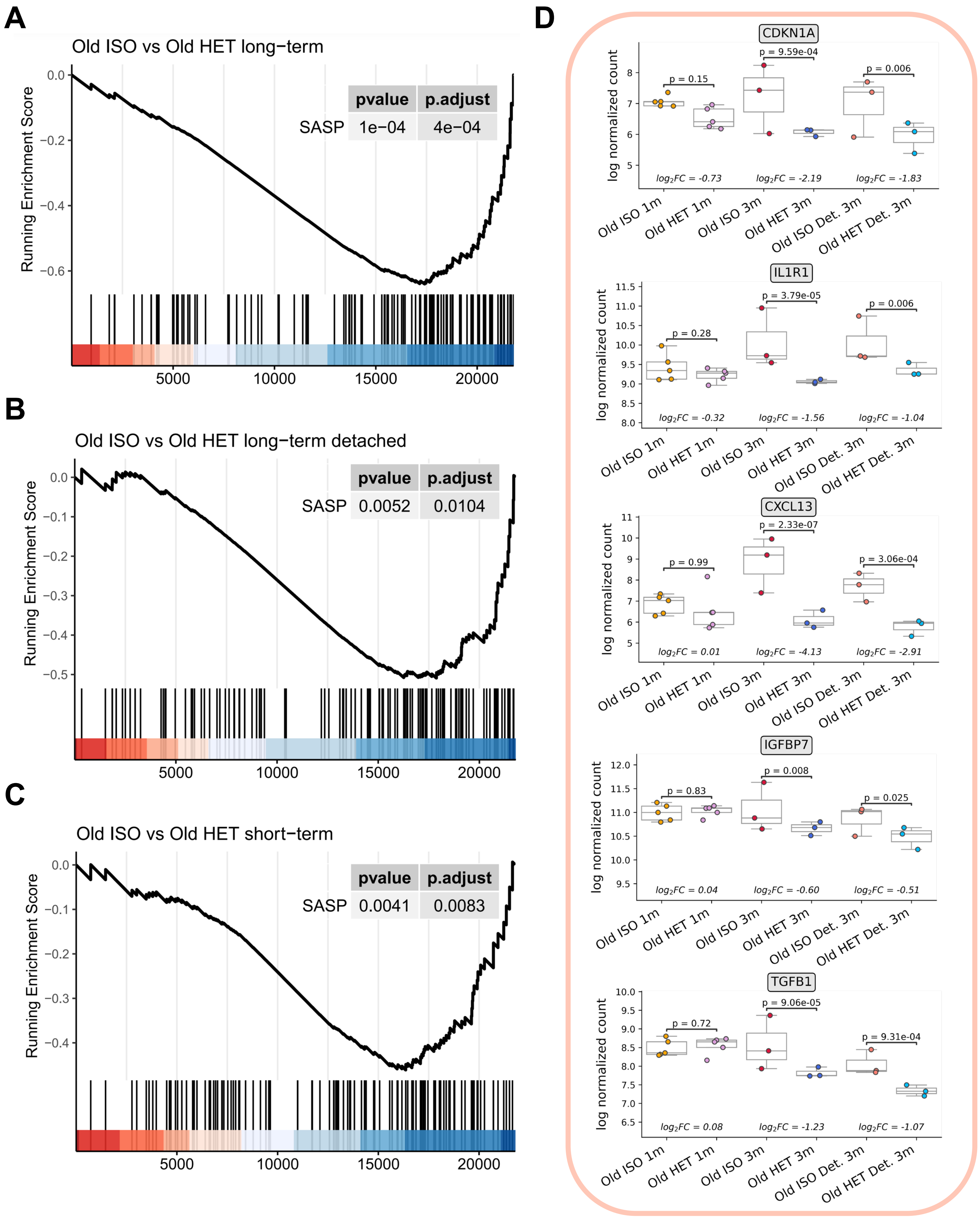
